# SVCROWS: A User-Defined Tool for Interpreting Significant Structural Variants in Heterogeneous Datasets

**DOI:** 10.1101/2025.01.24.634734

**Authors:** Noah Brown, Charles Danis, Vazira Ahmedjanova, Jennifer L. Guler

## Abstract

Genomic structural variants (SVs) are pervasive and can impose major phenotypic impacts. However, it is difficult to appreciate the individual significance of SVs when they are heterogeneously positioned across a genomic neighborhood. Further, ubiquitous variance in SV calling accuracy complicates SV counting and downstream analysis. Tools exist to simplify SV datasets but they are not suited for all applications. Here, we present a new SV merger, SVCROWS: Structural Variation Consensus with Reciprocal Overlap and Weighted Sizes. This option-rich merger summarizes SV regions using a size-weighted reciprocal overlap framework, accounting for skewed impacts of variable-length SVs. User input directs stringency, enabling various levels of resolution in complex genome regions that harbor a spectrum of SV sizes. Further, by optimizing SVCROWS parameters, the user can tailor results to their study system. When compared to other SV merging programs, SVCROWS maintained accuracy and conserved rare genotypes from both simulated and real-world datasets. Visualization of merger output was critical for identifying how some algorithms derived erroneous conclusions while SVCROWS remained reliable, especially in complex regions. Overall, the novel SVCROWS algorithm presents an improved framework for SV interpretation; its intuitive nature and generalizability facilitates its application to virtually any workflow.

**Graphical Abstract:** SVCROWS (Structural Variation Consensus with Reciprocal Overlap and Weighted Sizes) is a structural variant merger that leverages option-rich, size-weighted comparisons to better resolve complex inputs and separate out regions of meaningful biological differences.

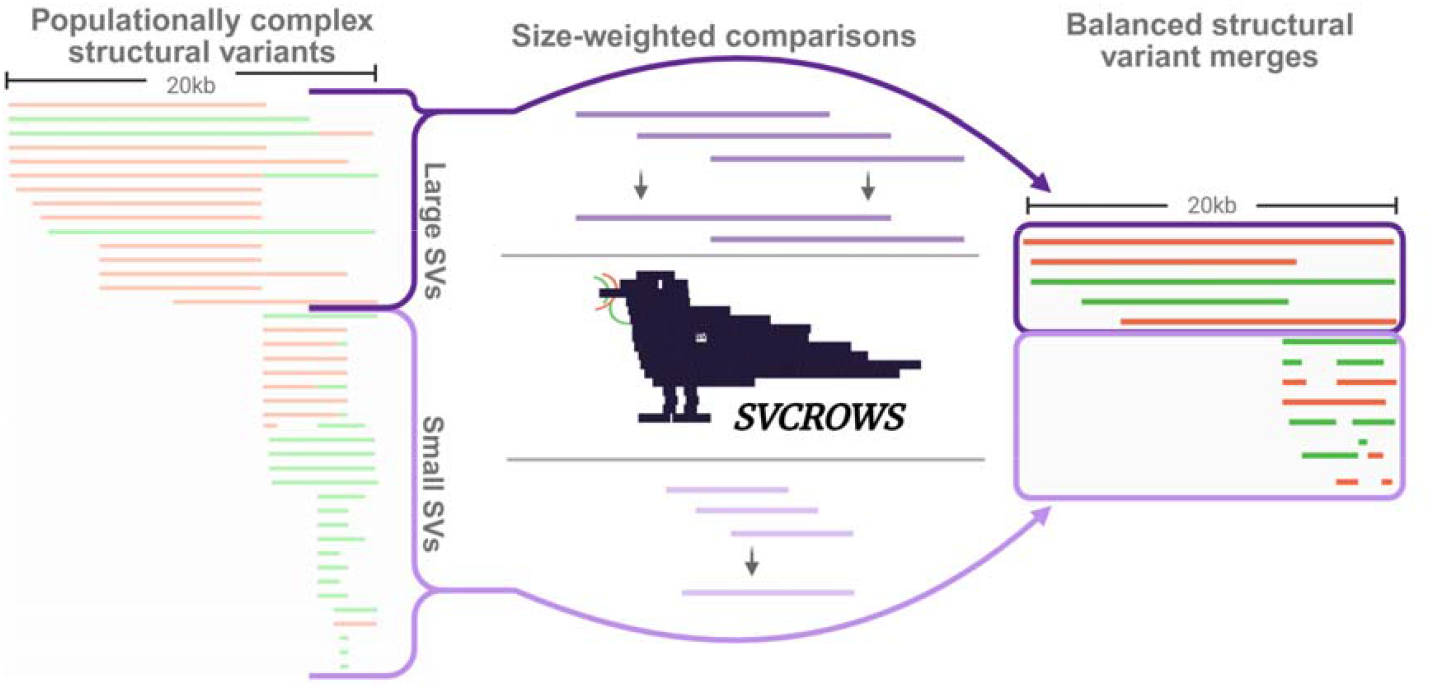

## Introduction

Genomic structural variations (SVs) are large (>50 bp) duplications, deletions, inversions, insertions, and/or translocations. SVs are both evolutionarily important for organisms and causative factors in many human diseases(1–5). Their abundance in all life makes them prime targets of study. Recent advances in sequencing technologies have improved detection of SVs, as well as the sequence accuracy and breakpoint mapping of SV calls(6). Validation of these programs and protocols has become more accessible using benchmarking datasets such as the *Genome in a Bottle* consortium(7), and the *1000 Genomes Project* consortium(8). However, newly developed tools and benchmarking programs largely focus on single-sample SV detection (i.e. from bulk sequencing), abstaining from comparisons across individuals in a population as is required for single-cell (SC) genomics.

As work determining the connection between rare genotypes and phenotypes becomes more common, researchers must evaluate more data with increased precision. Overlapping yet distinct SV regions arise either from biological variance or technical noise and make subsequent analysis more difficult. Therefore, accurate SV merging to reduce complexity is a key step prior to SV interpretation which can involve either checking for an expected genotype (i.e. validation) or detecting new genotypes (i.e. discovery). Thus, the outcome of SV merging is an influential step in analysis pipelines. In the case of both validation and discovery, merged SVs have greater statistical strength to support biological conclusions(6, 9–12). Determining how to merge SVs is challenging because small shifts in position can have major impacts on SV interpretation. SV merging must maintain a balance between removing redundancy across a dataset by merging similar SVs and preserving heterogeneous and rare SVs.

The best way to address these merging challenges remains debated, but strides have been made by several SV merging algorithms to reconcile this issue(10, 13–19). However, these existing approaches have limitations. First, many rely on high-accuracy inputs and have only been validated in that context. This is particularly an issue when sequencing depth is limited (e.g. for SC genomics), the reference genome is low quality, or SVs sit in highly repetitive regions of the genome. Large population studies also remain difficult to resolve despite high sequencing quality, because the increased SV heterogeneity leads to overmerging(14, 18, 20). Another common limitation is incompatibility of mergers with different data types, because many mergers are incorporated into larger analysis pipelines to increase effectiveness. This places constraints on the input type or upstream SV caller, invalidating certain study designs. Further, while SV mergers generally have options to control SV matching parameters, their direct impact is not immediately transparent to the user without performing many iterations. Overall, under- or overmerging of variable regions across many samples leads to a loss of information or generates misleading information.

Here, we introduce a new SV merger, ‘SVCROWS’ (Structural Variation Consensus with Reciprocal Overlap and Weighted Sizes), that seeks to overcome these limitations. SVCROWS merges SVs called by upstream methods (or any genomic intervals of interest) into representative SV regions (SVRs), so that technical variation is simplified while preserving rare alleles. This is accomplished through size-weighted reciprocal overlap (RO) comparisons that adjust the stringency of merging in response to SV size. Here, we demonstrate that size-weighted comparisons enable SVCROWS to achieve higher performance in both simulated and real-world SV datasets, particularly in complex genomic regions. These results can be further improved by parameter optimization and selection of an advantageous run mode. SVCROWS also benefits from a focus on biologically oriented features and compatibility with both up- and downstream steps of genomics protocols.

## Materials and Methods

### A framework of size-weighted comparisons to capture significant SVs

SVCROWS uses the principle of reciprocal overlap (RO) thresholds to determine whether to place SVs in the same SVR (i.e. matching/merging SVs). While the RO-based approach is used in existing applications(19, 21, 22), these rely on a single, fixed RO threshold value. This static method overlooks the disparity in SV lengths and their potential biological impacts, creating a false equivalency between small and large SVs. For instance, if we measure biological impact by the number of genes within an SV (**Fig. 1A-B**), a larger SV is more likely to have a greater effect (e.g. encompasses more genes and regulatory regions) on the phenotype than a smaller SV (e.g. encompasses few genes or even a promoter region). Specifically, as depicted in **Fig. 1A-B**, there are two obvious inconsistencies in treating different sized SVs with the same stringency: 1) SV2 matches both large and small reference SVs, despite causing a 3X increase in potential impact in the larger SV (i.e., involving 3 genes); 2) When either SV size lacks just one gene compared to the reference (i.e. the same level of biological difference to the reference), the small SV matches at a 55% RO (**Fig. 1A**, SV2) while the larger SV matches at 95% (**Fig. 1B**, SV1). Using this reasoning, we posit that when a size-static matching parameter like RO is used, it is more likely to reflect genotypes disproportionate to their impact. This is the basis for the size-weighted comparisons used by SVCROWS.

**Figure 1.**
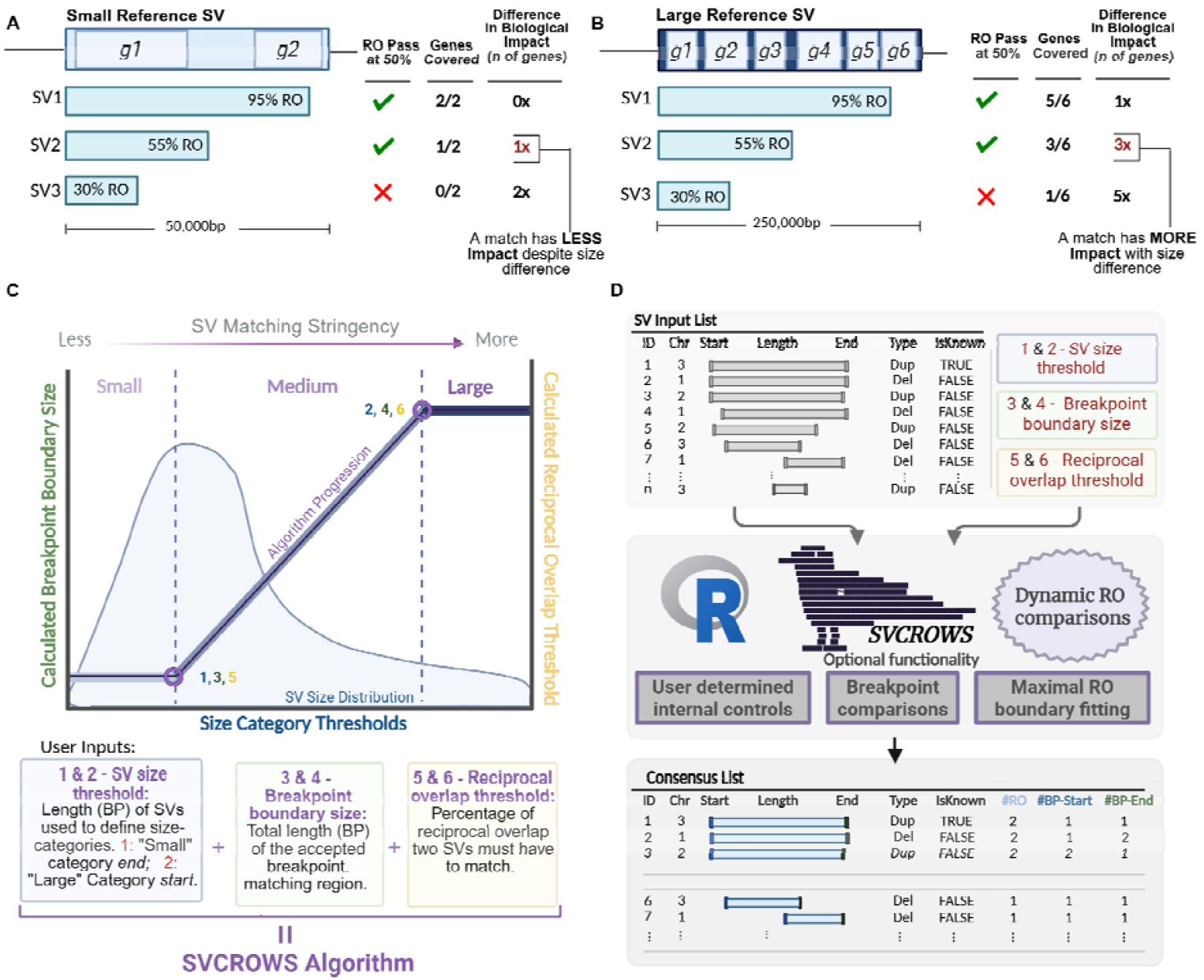
The SVCROWS algorithm compares SVs with regard to size, and potential impact. **A)** A small (50,000bp) reference SV (light blue) encompassing 2 genes and, **B**) A large (250,000bp) reference SV (dark blue) encompassing 6 genes; being compared to 3 potential SVs (teal) at a static 50% Reciprocal Overlap threshold. Using each gene as a unit of biological impact; smaller SVs are more tolerant to changes in size when reflecting similarity to the original SV. **C**) Six inputs defined by the user weight the reciprocal overlap determination by size. x-axis: “Small” and “Large” SV category thresholds (variables 1&2). Left y-axis: The width (in base pairs) of BP-Matching region (see **Fig. S1**, variables 3&4 respectively). Right y-axis: The required level of RO-threshold (as a percentage) for SV matching (variables 3&4 respectively). These inputs generate the SVCROWS Calculation Trajectory (purple) which defines the dynamic RO stringencies. **D**) Logical workflow of SVCROWS merging function. A list of SVRs is produced, from a single input list. The number of matches is tabulated (bottom). ‘Dup’ = duplication, ‘Del’ = Deletion, ‘#RO’ = Number of matching SVs, ‘BP-Start/End’ = matching breakpoints on the 3’/5’ ends of the SVs. Options to augment the run mode are also available.

### Selecting parameters to dictate stringency using size-weighted comparisons

To address high variance in SV length and position, the user directs SVCROWS to increase or decrease the requirements of the RO threshold and other factors involved in the decision to merge two SVs (i.e. stringency) according to the SV size (**Fig. 1C**). Dynamic RO thresholding by SVCROWS involves 6 input parameters to define the stringency based on analysis needs (**Fig. 1C**, colored boxes). First, SVCROWS creates size categories to group SV-comparison stringencies by SV size. By default, these parameters are determined by the first and third quartiles of SV length in the given dataset. These can be further refined based on the minimum and maximum SV sizes the user considers relevant in their organism or study interest. For example, in an analysis of CNVs, the ‘small’ category could be the average exon size while the ‘large’ category could reflect the average size of CNVs reported in that genome (**Fig. 1C**, blue inputs 1 & 2). Second, users dictate bounds that allow SVCROWS to match breakpoints between input SVs (**Fig. 1C**, green inputs 3 & 4, **Fig. 1A**). Finally, users define RO thresholds at the boundaries of the size categories (**Fig. 1C**, yellow inputs 5 & 6). Collectively, these input parameters set the SVCROWS algorithm to respond to SV size (**Fig. 1C** red line). Detailed parameter definitions and functions are provided in the ***Supplemental Methods***.

### SVR consensus generation

SVCROWS merges SVs and organizes data into a table of distinct SVRs according to the input parameters. The algorithm processes SVRs in descending order of length (**Fig. 1D, top**). Therefore, the largest SV in the region dictates boundaries, and the breakpoints reflect those provided by the upstream SV-caller. When used as recommended, the stringency is higher for larger SVs than smaller SVs (**Fig 1D**); however stringency can be assigned so that the opposite is true as well. SVCROWS quantifies matching SVs on the number of RO and SV breakpoint (BP) matches at 5⍰ and 3⍰ ends of an SV (**Fig. 1D**, bottom).

### Optional inputs and functions

In addition to size-weighted merging (“Scavenge” mode), SVCROWS incorporates several features to construct and quantify SVRs. While some features mirror existing functionality, SVCROWS integrates these functions into a single platform and introduces novel functions (denoted with * below). Seven major optional features are outlined below, with detailed descriptions available in the ***Supplemental Methods:***

#### i. *Automatic input parameter calculation

SVCROWS calculates six input parameters based on the first and third quartiles of input SV sizes. This allows SVCROWS to dynamically adjust to the specific characteristics of the dataset.

#### ii. *Breakpoints as a secondary SV matching factor

Users can opt to incorporate breakpoint matching to aid SV merging (**Fig. S1A**). When enabled, SVCROWS reduces the required RO threshold to its minimum value (user input 5) for SVs with matching breakpoints. This feature is useful for identifying matches in repetitive genomic regions where one breakpoint is consistent, but the opposite end is misaligned.

#### iii. *Use of ‘Known’ regions in input

A user can define input ‘known’ SVs, which reduces all requirements for matching in that region to only a single base pair (**Fig. S1B**). For example, by defining the region of *AMY1* in the human genome as a ‘known’ CNV(23), a user can collapse all variation in this genomic neighborhood.

#### iv. *Convenient annotation of all input SVs by sample

SVCROWS uses a custom input format that allows the program to individually track the outcome of all input SVs. This is especially convenient for workflows that require per-sample analyses.

#### v. Expansion of SVRs to maximum size

The user can employ this option to set SVR size based on the maximum and minimum matching breakpoint of all SVRs in a region (**Fig. S1C**). This functionality may be useful when downstream applications benefit from larger SVRs (e.g., gene ontology studies).

#### vi. Querying against a list of features

The user can alternatively use the “Hunt” function, which takes a secondary input - a ‘feature list’ - for comparison with an input list of SVs (or SVRs). For example, this functionality allows the user to compare a set of genes to SVRs determined from “Scavenge mode” using the size-weighted framework.

#### vii. Enumeration of other variables

As SVs match, the user can provide numeric information that 1) takes the number of reads originally used to call the SV and adds them together and 2) takes the quality score of each SV and averages them together.

### SV merging for SV-Simulation, SC dataset, and HG002 Truth Set

In order to understand how different sequence-independent merging algorithms perform over diverse datasets, we compared SVCROWS to several SV mergers with a variety of algorithm types: Jasmine(17), Survivor(24), SVimmer(15), Truvari(18), and CNVRuler(19). We chose these five SV mergers because they are compatible with most upstream SV callers, and present different strengths and weaknesses in their composition. To normalize for these differences, we instructed each program to function with parameters as close to the default parameters of SVCROWS-Default as possible; using the ‘default’ parameter assignment with breakpoint matching and RO expansion options enabled (**Fig. S1**). Default SVCROWS parameters use the first and third quartiles of SV sizes to demarcate size boundaries, 10% of respective lengths to set the size of breakpoint boundaries, and 40% and 70% as the small and large RO thresholds. Unless otherwise indicated, SVCROWS is run in ‘default’ mode, which uses both the breakpoint matching (function ii) and expand-RO (function v) features.

The precision of a typical CNV calling program ranges from a single basepair to no more than 500bp. Because all mergers (excluding SVCROWS) include breakpoint distance to some extent in merging decisions, we ran all mergers with a lenient 1000bp buffer region around either end of the SV; this is larger than previously used in order to encourage SV comparison(15). See ***Supplemental Methods*** for the precise command-line run for each program.

We manually compiled the SV input lists for SVCROWS and CNVRuler and organized them into TSV files used for each run matching their individual formats (**Tables S2 & S3**). Jasmine, Survivor, and SVimmer accept the raw VCF files from the SV caller. To normalize for any inconsistencies between output from each caller, all alternative alleles were manually converted to either “<DUP>” or “<DEL>“, and we replaced the sample field with a “.”. SVs were first consolidated for Truvari as recommended in their user wiki BCFTools(25), which essentially concatenates all SVs together without merging. We noted this concatenation step with BCFTools led to artifacts in complex regions in our dataset (see **Fig. S10B**) and opted to use a different BCFTools command that merged purely redundant SVs (i.e. those with precisely matching breakpoints) before further merging. This largely fixed the artifacts (See ***Supplemental Methods*** for more details). Truvari notably has the optional ability to use sequence information contained within aligned reads, which may assist in refining some genotype calls. However, because SVCROWS and other mergers cannot incorporate this information, and SVCROWS is intended to be generalizable to any workflow, we used Truvari’s otherwise robust merging algorithm without this option.

### SV-Simulation and analysis

To effectively test the capabilities of SVCROWS, we developed a custom script that simulates SVs as they appear in datasets with many samples, such as SC datasets. Using this SV simulation, we generated a simulated-Truth Set SV list and subsequently derived per-cell observed SVs that incorporate both biological heterogeneity and technical noise that are propagated to simulated individuals (‘cells’). The command-line interface (CLI) accepts several inputs that control key dataset characteristics and is fully described in the ***Supplemental Methods***. Briefly, we divide the simulated-Truth Set SVs into ‘frequent’ events (prevalence >80% of cells; referred to as ‘clonal’ in the CLI) and ‘infrequent’ events (events with frequencies drawn from a geometric distribution to mimic stochastic occurrence in one or a few cells; referred to as ‘subclonal’ in the CLI). The simulator randomly determines the population frequencies of common and rare SVs before constructing cells, ensuring that each simulated cell matches those frequencies (e.g., if a feature occurs at 10% frequency in 100 total cells, the simulator assigns it to 10 randomly chosen cells). The simulator samples SV coordinates across a region of defined size (this study used the length of HG38 chromosome 1) with log-uniformly distributed lengths, allowing overlaps. To mimic genomic instability, the simulator generates high-complexity regions that guarantee multiple simulated-Truth Set SVs overlap within a defined locus. Finally, to simulate real-world uncertainties in SV calling, the simulator introduces positional and size fluctuations to per-cell SVs. A single variance parameter controls the fluctuation magnitude: low values preserve true coordinates, whereas high values allow large shifts (up to ∼30% positional offset and 70% size distortion). Larger CNVs experience proportionally greater absolute deviations (see **Fig. S2** for an example run of the simulation).

To analyze the SVR output of the simulation, we compiled the features divided among cells to match the input format required by the merger tools (Jasmine, Truvari, and SVCROWS-Default). We then compared the resulting outputs to the simulated-Truth Set using Bedtools intersect under three static reciprocal overlap thresholds (50%, 80%, and 95%). We counted true positive (uniquely matched against the simulated-Truth Set), false negatives (non-unique matches to the simulated-Truth Set), and false positives (SVRs without matches to the simulated-Truth Set). We used these values to calculate the F1 score (see **Fig. 2A** and ***Supplemental Methods***).

**Figure 2.**
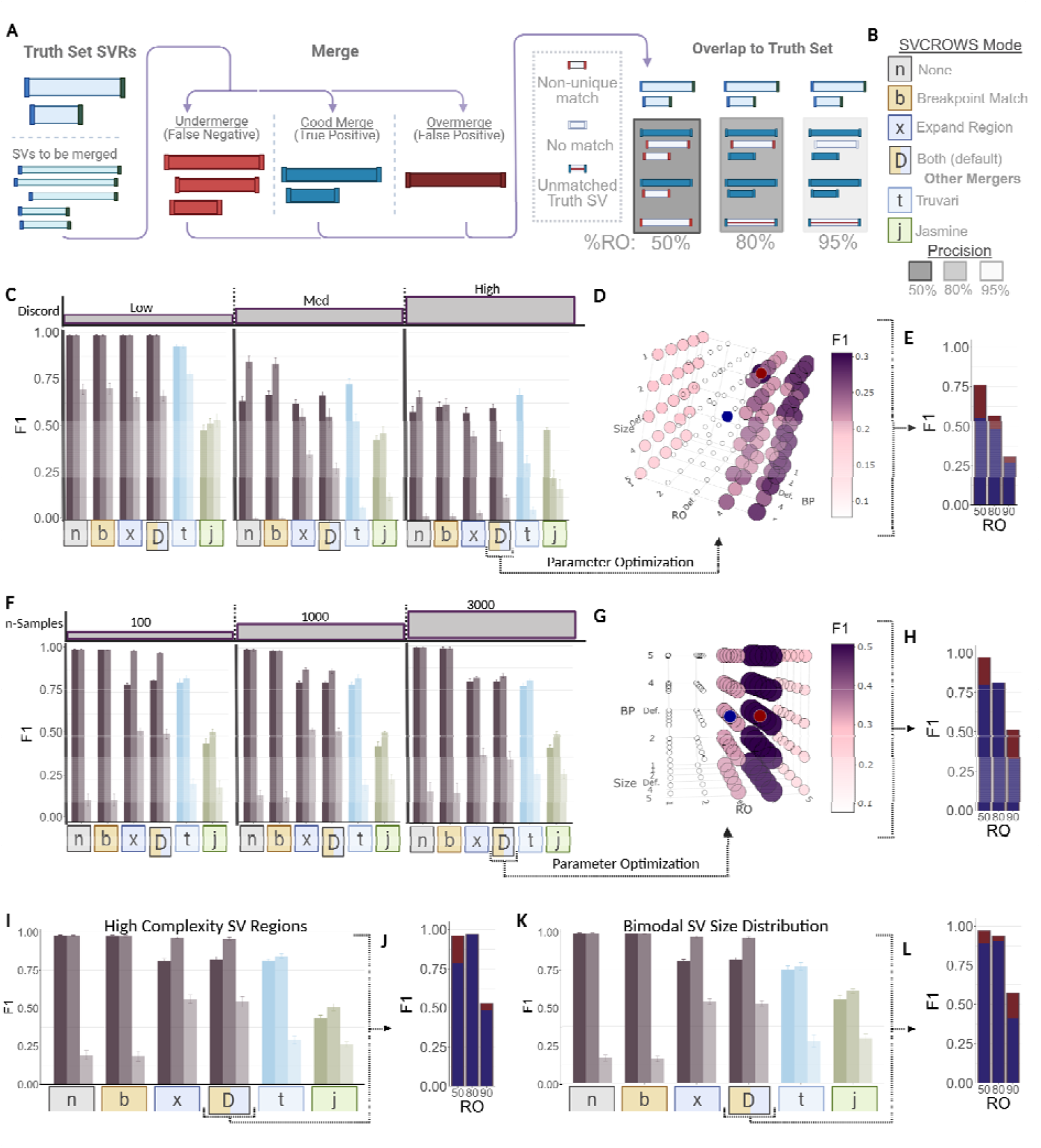
Optimization of SVCROWS input parameters compared to other SV mergers on simulated datasets. **A)** Diagram illustrating how merger accuracy and precision are determined across three levels of stringency relative to the simulated-Truth Set. **B)** Key for mergers and options used for comparisons on simulated SV data (applies to subsequent figures). **C-L)** Comparison and optimization of F1 scores in four challenging simulations. Each merger or option set depicts three bars at varying levels of reciprocal-overlap stringency; each set of bars represents the same post-merge dataset. Error bars indicate the standard deviation of the F1 score. Each simulation tests one aspect of input variance while keeping others constant: **C-E)** Three levels of discordancy (fluctuations in start, end, and size of SVs). **F-H)** Three levels of sample number (100, 1000, 3000). **I-J**) High-complexity regions containing several overlapping SVs of various sizes and frequencies. **K-L)** A bimodal distribution of SV sizes in which very large SVs are interleaved with ∼100 × smaller SVs. **D, G)** Representative 3D visualizations of F1 scores from a single constituent SVCROWS-Default replicate (one of five replicates), showing systematic adjustment of the six input parameters required at 95% RO relative to the simulated-Truth Set. The **blue** dot represents the unoptimized F1 score at 95% RO, and the **red** dot represents the highest F1 score after parameter optimization (for all 3D plots, see **Fig. S4). E, H, J, L)** Representative graphical results of parameter optimization for the same replicate. The **blue** bar indicates the original F1 score, and the **red + blue** bar indicates the increase after optimization.

### SVCROWS parameter optimization using simulated data

To highlight the relationship between SVCROWS input parameters, we employed a 3-dimensional analysis that allowed us to visualize the interplay between the three input axes (Size, RO, and BP) using the F1 score. For each selected dataset, we generated an interactive 3D scatter plot using the plotly(26) library in R. The x-, y-, and z-axes correspond to Size, RO, and BP parameter indices, respectively, while color and size indicate F1 score. For each of the selected simulated SV runs where this 3D analysis was performed, we produced a series of 5 levels of stringency based on the predetermined SVCROWS-Default values (see ***Supplemental Methods*** and **Table S1** for full specification). Ultimately, this visualization enables rapid identification of optimal parameter combinations where F1 scores are maximized, thereby facilitating comparative evaluation of algorithmic performance across the full 5×5×5 parameter space.

### Single Cell dataset acquisition and SV calling

We used pyega3 to download the single-cell ovarian cancer dataset, EGAD00001009455, with permission of the European Genome Archive (EGA)(27). The authors note their compliance with best practices detailed by the EGA. The dataset consists of 326 single cell (SC) whole genome sequences derived from a solid ovarian cancer tumor from a single patient (termed SC dataset). Notably the patient also had triple negative breast cancer. The full Galaxy-based pipeline can be found at https://usegalaxy.eu/u/sillycrow/w/lumpy. Briefly, we used downloaded BAM files to generate files containing only split and discordant reads as determined by previous sequence alignments. Then, all three BAM files are used as input into the LUMPY SV caller. We chose LUMPY for this analysis due to its high precision breakpoints and utility in single-cell datasets(28–30). We used default parameters except for the expected read length (150 base pairs) and the minimum read mapping quality for inclusion (≥5). For this analysis, we excluded BAM files with less than 1000 elements in either the split-read or discordant-read files due to issues with LUMPY performance in low-read support samples(31, 32). Additionally, we limited the SC analysis to only duplications and deletions with a length of <1Mb called by LUMPY for simplicity. Prior analysis of this dataset detected high frequency SVs with variable breakpoints, a common feature in cancers(33). Overall, 4771 total SVs were detected across the dataset (**Tables S2 & S3**). For comparison of SVCROWS RO stringency in complex regions (see **Fig. 3C**), we tuned SVCROWS to interpret differences in the human genome by using the average size of a human exon (∼1.3kb(34)) and the average size of a gene-plus-intergenic region (∼38kb(35–37)) as small and large SV sizes, respectively.

**Figure 3.**
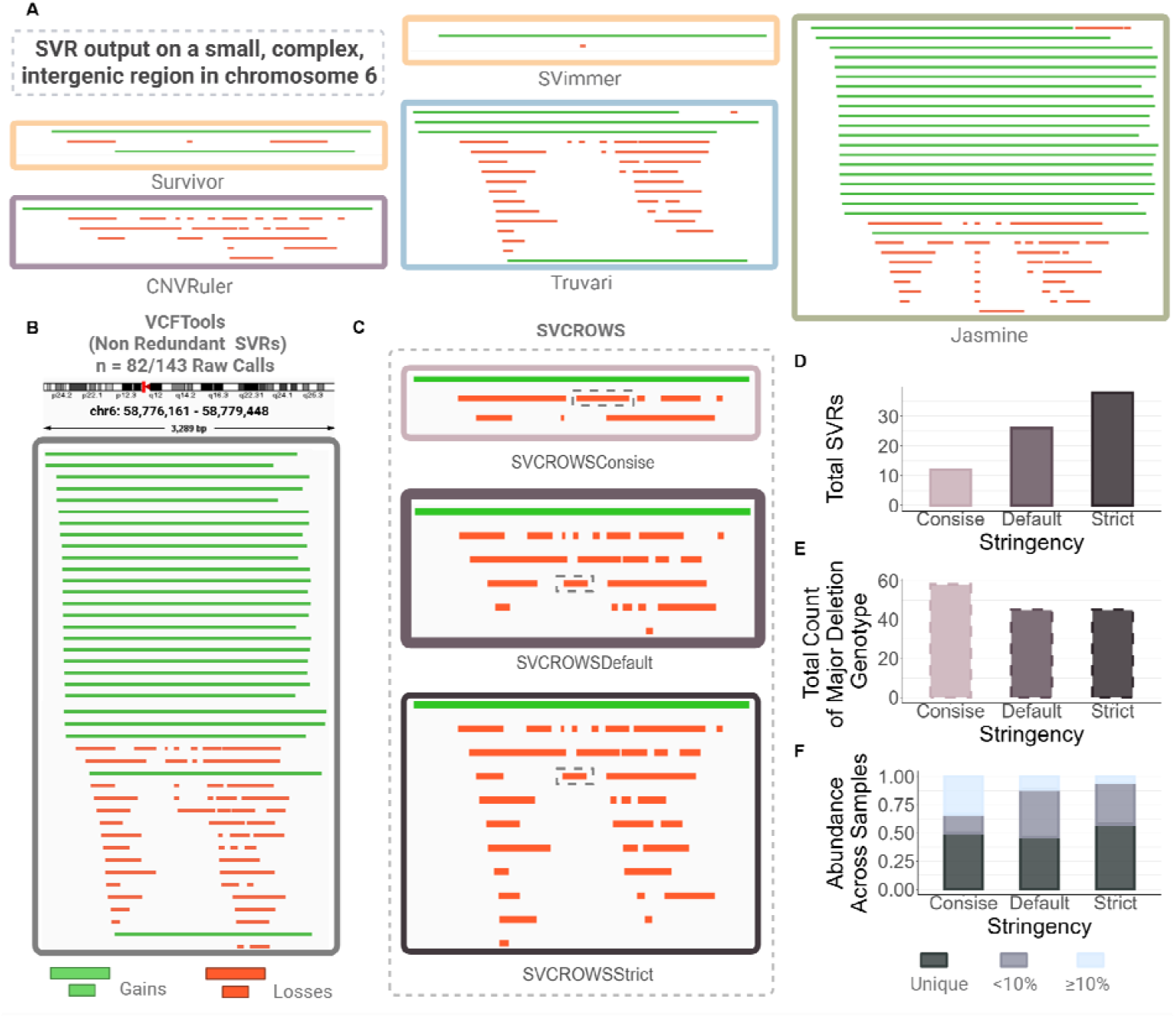
Options to control stringency and SVR construction in SVCROWS allows for optimized merging in a complex region. A single cell (SC) dataset consisting of 326 cells was used to evaluate SVR generation by multiple mergers. **A)** Integrated Genome Viewer (IGV) frames showing the output from the 5 other mergers. **B)** IGV view of VCFTools as a non-merged negative control. **C-F)** SVCROWS applied to the dataset over optimized stringencies (strict, concise, or default, see *Supplemental Methods* for more details): **C)** IGV view of resulting SVRs in a representative 3.2kb centromeric region of chromosome 6. Dashed boxes indicate the most abundant ‘loss’ genotype detected in the dataset. **D-F)** Quantification of panel **C. D)** Total number SVRs in the region after merging. **E)** Total number of SVs matched to the SVR representing the abundant deletion present in ∼14% of the population. **F)** Abundance of each of resulting SVRs across each merger. ‘Unique’ are those that only appeared once in the population, ‘<10%’ represent SVRs rare but present in more than one sample, ‘≥10%’ are common SVRs.

### HG002 dataset processing and analysis

The HG002-Truth Set (HG002_SVs_Tier1_v0.6.tsv) is part of the *Genome in a Bottle* (GIAB) consortium, a high confidence, multi-validation genome assembly(38–42). This dataset consists of 9641 individual SVs (>50bp) confirmed by at least 3 independent SV calling programs, including SV calls from both long- and short-read data. For this analysis, we used only Tier1 (GIAB certified highest confidence) SVs, including representative SVs that allow assessment of merger efficacy in regions with multiple SV calls across the same region. The HG002-Truth Set includes both deletions and insertions, but because not every merger or SV caller can handle insertions without the inclusion of raw sequences, we only used deletions in this analysis.

To test SV mergers, we used HG002 SV calls from 5 different programs as input: LUMPY(30), pbsv(43), Sniffles(44), MrCaNaVaR(45), and MetaSV(46). We combined all calls into a single ‘Aggregate List’, many of which are not part of the HG002-Truth Set (∼90%), either because they are false positive calls, or they did not reach the threshold to be called in the Truth Set (i.e. called by 3 programs, pass various quality filters, etc.). Further, because the largest SV in the HG002-Truth Set was ∼1Mb, we excluded SVs >2Mb from the analysis. This represents the manual filtering step, which is part of most SV calling pipelines (see **Fig. S9** for a diagram of the HG002 dataset processing). We first calculated Jaccard indices of the SV mergers using the Bedtools-jaccard method on this cleaned Aggregate List, which considers both the coverage and depth of represented regions to calculate differences in merging (see ***Supplemental Methods*** and **Table S4**).

Many of the SVs called in the cleaned Aggregate List are perfect (or near-perfect) matches the HG002-Truth Set and were used as true positives in an effort to establish relative merger accuracy. To do this, we filtered the Aggregate List and HG002-Truth Set to make a reference set containing only SVs common to both. As such, we eliminated irrelevant false negatives and positives (i.e. those that are present in the Aggregate List but not in the HG002-Truth Set, or vice-versa) to only those SVs that had any overlapping bases ([Aggregate_List ⍰ HG002-Truth_Set] ≥ 1bp). The resulting pared-down reference set of SVs is dubbed the ‘On-Truth Set’ and used to compare the filtered Aggregate List against. This filtering ensured that only SVs present in both sources, hence theoretically recoverable by any merger, were evaluated as potential true positives. (see **Fig. S9**). The filtered Aggregate List is then merged and then compared to the On-Truth Set using F1 score in the same method as above (**Table S5**).

Because inclusion of an SV in the HG002-Truth Set depends on tight breakpoint concordance among multiple callers, even small positional variances can cause otherwise correct calls to be excluded. However, both accepted and rejected calls remain in the filtered Aggregate List. Therefore, this study design leveraged the technical variance and noise in the calls made by the 5 callers to assess merger’s capabilities to reproduce the On-Truth Set purely computationally (as opposed to the original, labor-intensive HG002-Truth Set construction). The recaptured similarity to the On-Truth Set was assessed across different overlap thresholds to highlight how faithfully breakpoints were conserved after merging the Aggregate List, we then overlapped the SVRs against the On-Truth Set to calculate true positives and F1 score as above.

## Results

### Optimization of SVCROWS for outstanding performance under challenging input conditions

To benchmark the performance of our new size-weighted comparison framework, we evaluated SVCROWS using different combinations of its optional decision-making functions (breakpoint matching and expand-RO; **Fig. S1**) alongside two widely used SV mergers, Truvari(18) and Jasmine(17), in a controlled SV-simulation environment. We used a custom script to generate SV input lists with precise control over key characteristics, including event size, breakpoint variance (“discordancy”), local complexity (multiple overlapping SVs), and the number of simulated cells (**Fig. S2**; see ***Supplemental Methods***). These datasets were compared against a simulated-Truth Set to systematically evaluate sensitivity and precision under increasingly challenging input conditions (**Fig. 2A–B**). Importantly, the simulations reproduced expected patterns: F1 scores declined as input complexity increased, confirming that the benchmarking system captured realistic SV-merging challenges (**Fig. 2C, F**).

Comparisons were affected by trade-offs between false positives and false negatives when assessing similarity to the simulated-Truth Set (**Fig. 2A**). To address this, we measured merger performance using three reciprocal-overlap (RO) thresholds - 50%, 80%, and 95% - which emphasize recall, balanced accuracy, and precision, respectively. Across the four simulation scenarios (**Fig. 2**), SVCROWS consistently outperformed competing tools, even under challenging inputs. Under increasing discordancy, SVCROWS-Default retained higher precision than either Jasmine (F1 1.54× higher) or Truvari (F1 1.49× higher) across comparisons. Although breakpoint shifts reduced overall F1 scores (**Fig. 2C–E**), SVCROWS maintained superior performance. With escalating sample sizes that increased opportunities for merging errors, performance declined across all methods, yet SVCROWS-Default maintained a distinct advantage, with mean F1 scores 1.84× and 1.39× higher than Jasmine and Truvari, respectively (**Fig. 2F–H**).

In high-complexity regions containing many overlapping SVs, SVCROWS-Default produced far fewer false positives and false negatives than either Jasmine or Truvari (2.20× and 1.40× fewer, respectively), highlighting the utility of its size-weighted comparisons. This corresponded to a increase in F1 at the 95% overlap threshold - 2.06× and 1.86× higher than Jasmine and Truvari, respectively - in crowded genomic contexts (**Fig. 2I, J**). Finally, in datasets with bimodal size distributions, where very large SVs co-occurred with much smaller events, SVCROWS-Default showed a superior balance of recall and precision, outperforming other methods across both liberal and stringent overlap thresholds despite its reliance on size distribution to determine stringency (**Fig. 2K-L**).

The trade-off for SVCROWS’ increased accuracy and precision is computational time. Because SVCROWS must perform comparisons for each SV individually, small input lists execute almost instantly, whereas increasingly larger datasets require proportionally more time than Truvari or Jasmine (**Fig. S3**).

Further optimization of SVCROWS input parameters substantially improved performance beyond the SVCROWS-Default run mode. Systematic tuning of breakpoint, size, and overlap factors revealed strong interactions among parameters, making multidimensional exploration essential (**Fig. S4**). Three-dimensional visualizations of the parameter space showed that optimized settings produced F1 scores exceeding the defaults at both 50% and 95% overlap thresholds (**Fig. 2D, G**). For example, under discordant and high complexity conditions, parameter optimization elevated F1 scores well above those achievable by Jasmine or Truvari at 95% RO, while simultaneously improving recall at 50% RO.

There were noticeable advantages among the distinct combinations of the four SVCROWS run modes. For example, under high-discordancy inputs, the use of breakpoint matching was preferable to expand-RO alone (**Fig. 2C**). Expand-RO by itself offered an advantage in almost all scenarios, while combining both functions helped offset the biases of each.

Together, these results demonstrate that the size-weighted framework implemented in SVCROWS not only performs robustly across diverse and challenging SV scenarios but also benefits from systematic parameter optimization. The ability to tune parameters allows SVCROWS to achieve superior accuracy and precision compared to existing tools, particularly under conditions where SV heterogeneity and breakpoint uncertainty complicate merging.

### SVCROWS allows control of SV merging in a complex single cell dataset

While SVCROWS performs well compared to other merging programs in simulated data across all tested scenarios (**Fig. 2**), it proves especially proficient at handling highly discordant and high complexity regions (**Fig. 2C** and **2I**, respectively). However, simulated data cannot recapitulate all forms of natural and technical variance found in real-world data. As such, we sought to further evaluate the performance of SVCROWS using SVs called from complex regions in a SC cancer dataset.

We compared SVCROWS’ size-weighted algorithm to the performance from 4 major types of SV mergers including: distance-based algorithms (SVimmer(15) and Survivor(24)), a minimum spanning forest-based algorithm (Jasmine(17)), a multifactored algorithm (Truvari(18)), a static RO algorithm (CNVRuler(19)), as well as a pure redundancy removal algorithm (VCFTools(47)) denoted by color below (**Fig. 3A, B**). Across the SC dataset, two regions stood out in complexity (>10x more SVs per 10kb than the background) including one complex sub-centromeric region on chromosome 6 (**Fig.3**), and a second large intergenic region on chromosome 10 (**Fig. S5, 6**). We focused our initial assessments on these regions to assess merging patterns across programs.

In the chromosome 6 region, there were 143 unmerged SV calls, with only 43% harboring breakpoints that exactly matched another SV (i.e. pure redundancy). Using only pure redundancy to define like-SVs is inadequate to fully simplify the region; evidenced by the undermerging of the ‘gain’ SV present in the majority of samples (**Fig. 3B**). After merging of pure redundancy, 82 SVs remained. When we visualized how the merging algorithms perform, it was apparent that they prioritize different aspects of SVs for merging (**Fig. 3A, S5, S6**). For example, Jasmine undermerged both small and large SVs across this region (**Fig. 3A**, green). Overall, in contrast to its satisfactory performance on the highly discordant simulated dataset (**Fig. 2C**), Jasmine failed to reduce complexity of this region compared to other programs, perhaps due to the large range of SV sizes. Conversely, distance-based algorithms seemed prone to overmerging where output had removed key differences (**Fig. 3A**, yellow). Truvari likely undermerged smaller SVs, leaving overlapping SVs with as little difference as 90bp (9.5% of the total size) unmerged. Further, by comparing SVCROWS to CNVRuler, we observed that the size-weighted RO led to substantial changes in the outcome of large regions. These observations are largely consistent with the chromosome 10 region (**Fig. S5, 6**). In both cases CNVRuler output only appears similar to SVCROWS but overall makes several repeatable errors in complex regions, that lower confidence in its consistency (highlighted in **Fig. S7**).

Observations in these specific regions agree with those made when broadening the scope to the entire SC dataset consisting of 4771 SVs (**Fig. S8, Table S3**). Specifically, we detected a wide range of resulting SVR sizes (**Fig. S8A**) and total SVR numbers, that mirrored our observations from the complex region on chromosome 6 (**Fig. S8B**). Merger choice also impacted summarized frequencies of losses and gains (**Fig. S8C**). It is typical for studies to note the proportion of gains and losses, but depending on the merger used, a researcher might either report similar numbers (**Fig. S8C**, Jasmine) or conclude there is a bias towards deletions (**Fig. S8C**, SVimmer). As such, some biological conclusions may depend on merger choice.

To demonstrate the impact of stringency on SVR discovery across complex regions (**Fig. 3C, C-F**), we compared SVCROWS-Default to 2 other stringencies determined by biologically-relevant characteristics (see *Materials and Methods*). At all three levels, SVCROWS maintained the major genotypes (the large gain-SV and boxed loss-SV), preserving the same breakpoints called in the input dataset (**Fig. 3E**). SVCROWS also conserved unique SVs (only appearing in a single sample across the dataset) indicating avoidance of overmerging (**Fig. 3F**). Through this analysis, we demonstrate 3 different SVCROWS applications that could be employed by a user; all of which reduced the complexity in the region, while maintaining the major patterns of SV frequency.

### SVCROWS produces accurate and sensitive SVR calls compared to other SV mergers

Using the SC dataset, we showed the utility of SVCROWS for handling complex regions with the goal of discovering SVs. However, without a single-cell (SC) Truth Set for comparison, validation of the merged results is not possible. To evaluate accuracy, we employed the HG002 benchmarking dataset(42) from the Genome in a Bottle (GIAB) consortium, which is considered the gold standard for high-accuracy SV benchmarking because of the extensive evidence used in its creation. After SVs were called using multiple SV callers and sequencing technologies, the HG002-Truth Set was compiled and merged largely through manual curation to determine the most accurate breakpoints. Thus, merging decisions could be validated or rejected based on factors beyond a simple algorithmic framework. Because such manual scrutiny is not feasible in most analyses, we sought to test how well computational mergers could recapitulate the HG002-Truth Set construction.

To accomplish this, we merged the outputs of 5 SV callers used in generating the HG002-Truth Set. Each caller contained false-positive and false-negative SV calls (**Fig. 2A**) in addition to true positives relative to the HG002-Truth Set. The datasets were combined, cleaned, and filtered into a single input, termed the ‘Aggregate List,’ for direct comparison of mergers. The incompleteness and inaccuracy inherent in these five datasets (despite their high quality) mimic the technical variance and error present in most real-world data. Correcting and summarizing the variance through accurate merging is therefore a critical step for downstream analyses. The post-merged Aggregate Lists were compared between mergers to assess overall similarity, and then back to a subset of the HG002-Truth Set containing only SVs also present in the Aggregate List. This subset - termed the ‘On-Truth Set’ - allowed assessment of merger accuracy without the confounding effect of unmerged, irrelevant SVs (**Fig. 4**; a diagram of data preparation steps is provided in **Fig. S9**).

**Figure 4.**
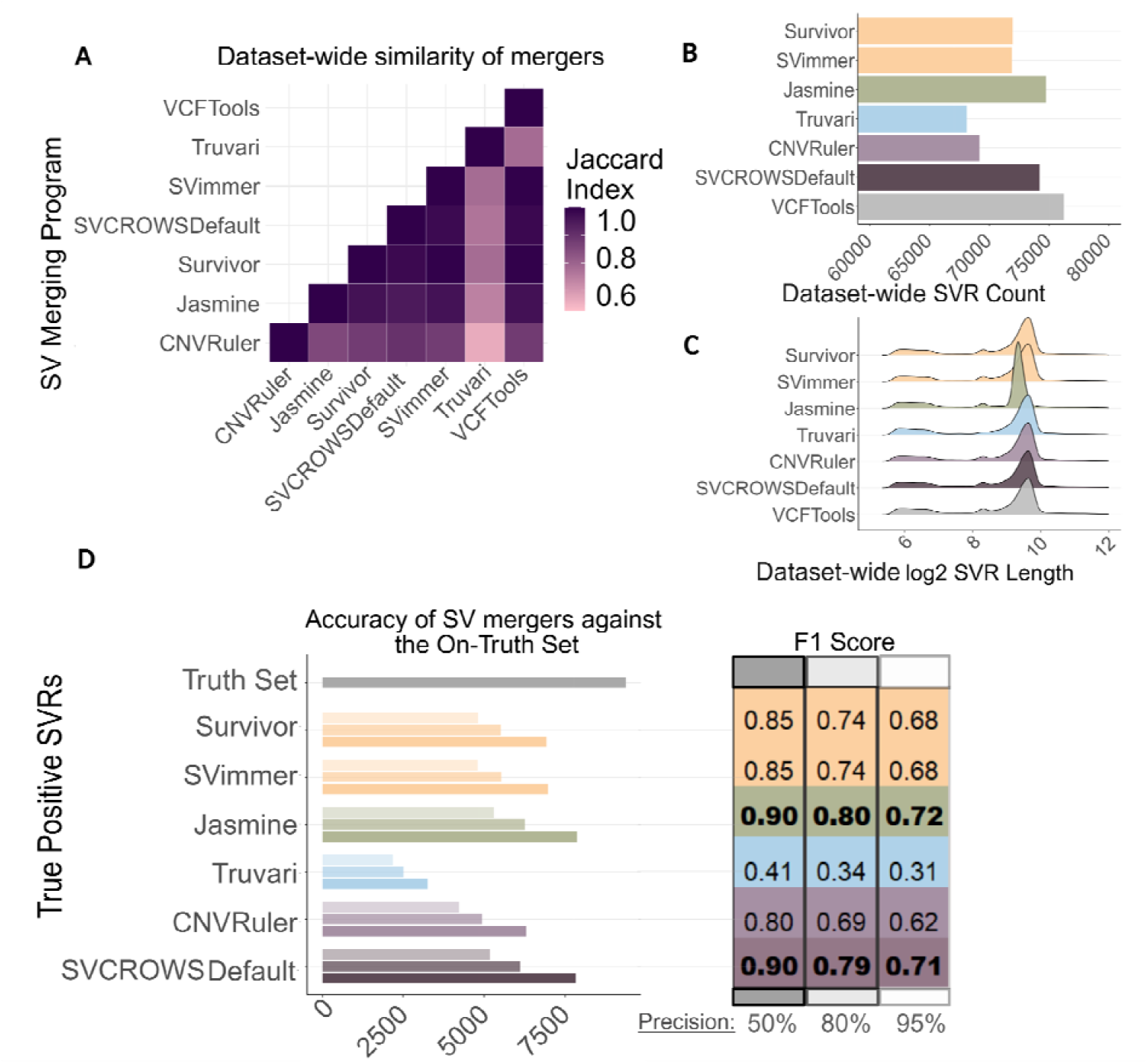
Comparison of SVRs from 6 different SV mergers to the HG002-Truth Set. We generated an SV Aggregate List from the GIAB consortium’s HG002 reference genome datasets consisting of calls from 5 different constituent SV callers. The Aggregate List was then merged using 5 different SV merging algorithm types (yellow; distance based (Survivor, SVimmer), green; minimum spanning forest based (Jasmine), blue; multifactored (Truvari), light purple; static RO (CNVRuler), dark purple; size weighted RO (SVCROWS), and gray; pure redundancy (VCFTools)), and then assessed for a variety of metrics. **A**) The Aggregate List was used as input to calculate differences in merging on a base-pair coverage level, and resultant Jaccard Indices for each merger are indicated (**Table S4**). **B**) Comparison of total SVR number after merging. C) Distribution of SVR sizes after merging. **D**) The resulting SVRs from above were then overlapped to a subset of the HG002-Truth Set (‘On-Truth Set’, see *Materials and Methods*) at 3 stringencies. Left) Accuracy of each merger as a rate of true positive calling. Right) Precision and Recall of merger performance represented by F1 scores using the following definitions: True positive SVRs uniquely overlapped to a single Truth Set SV. False negative SVRs exhibit non-unique matching or do not match to any Truth Set SV. False positives are those that had no overlaps to the HG002-Truth Set (**Table S5**).

The HG002-Truth Set has a relatively low density of SVs (only ∼2.5 SVs per 100-kilobases), but includes regions of high complexity as well as those with sparse SVs (**Fig. S10**). Flexibility in handling both cases is a challenging task for mergers. To objectively measure SVR coverage regardless of the level of complexity, we first used Jaccard indices to compare SV merger similarity on a per-base level. Higher Jaccard values reflect more similarity between lists generated by two different tools. Because each SV merger had the same input data (i.e. the Aggregate List), even minor differences in the Jaccard index represent regional under- or overmerging. Post merging, it was evident that SVCROWS displayed higher than average similarity to the other mergers (SVCROWS = 0.91, study average = 0.86), the wide range (0.62 – 0.99) of indices observed is consistent with our previous results suggesting that merger choice has major impacts on resulting SVRs (**Fig. 4A, Table S4**). For example, Truvari was more dissimilar across all comparisons (index average = 0.70), and CNVRuler was relatively more similar to other mergers (index average = 0.81), but still less than the study average (0.86). Further, because CNVRuler and Truvari both reported far fewer total SVRs than the other mergers (**Fig. 4B**), one could reason that both programs simply overmerge SVs. However, the Jaccard index between Truvari and CNVRuler is the lowest of the analysis (0.62) indicating algorithm-specific differences independently increased the rate of merging by each algorithm. Meanwhile, the similar SVimmer and Survivor algorithms exhibited a high Jaccard index (0.997).

In totality, each SV merger produced between ∼70,000-80,000 SVRs from the Aggregate List (**Fig. 4B**) with Jasmine and SVCROWS yielding the most SVRs. When we compared the size distribution of resulting SVRs, all SV mergers produced a nearly identical SV size range except for Jasmine (**Fig. 4D**). Further investigation revealed that Jasmine introduced SVR size alterations in the form of both truncations and extensions not present in the unmerged Aggregate List (**Fig. S10A**). Despite its dependence on size to define performance, the SVCROWS algorithm reflected a size distribution like other SV mergers, indicating a lack of size-related bias.

We then compared the merged SVRs to the On-Truth Set to assess merging accuracy using the same method as above (**Fig. 1A**). Jasmine and SVCROWS had the highest F1 scores of any SV mergers (**Fig. 4D**). In all 3 overlap stringencies, the number of true positive SVRs were ∼30% higher for SVCROWS and Jasmine compared to other programs while false positives were 20-40% lower on average. SVCROWS and Jasmine had similar false positive rates (only 3% higher for SVCROWS, on average across all stringencies). However, Jasmine consistently had false negative merges whereas SVCROWS produced none (**Table S5**). A lack of false negatives reflected SVCROWS’ tendency to avoid undermerging especially when regions lack complexity (**Fig. 4D**) and therefore circumvents loss of statistical power in downstream analysis.

Interestingly, in a reversal of the relative accuracy in the simulated datasets (**Fig. 2C, H**). Truvari’s F1 score was the lowest in this study, which agrees with its low Jaccard indices (**Fig. 4A**). When we directly visualized Truvari’s SVRs, we observed occasional merging artifacts in sparse regions that increase the rate of false negatives (**Fig. S10B**, examples 1 & 2). Further, in complex SVRs, Truvari fails to preserve genotypes from the Aggregate List (**Fig. S10B**, example 3). While SVCROWS also makes errors in the examples given, they can be rationalized by the input parameters chosen, unlike Truvari and Jasmine (**Fig. S10**). Combined, these inconsistencies led to both under- and overmerging by the Truvari algorithm. Overall, using a high-quality dataset, SVCROWS strikes a balance between accuracy and sensitivity to maintain the potential for rare genotype discovery with no indication of SV size-related biases.

## Discussion

SV datasets experience variance from increasingly large datasets(48) or by using different SV-calling applications on the same dataset(49). Resolving SV locations relies on accurate sequencing, precise mapping, and suitable coverage across a genomic region of interest. Such standards are hard to achieve, especially across repetitive regions like telomeric, centromeric, rDNA arrays(50, 51), or when using low-input methods like SC sequencing(52, 53). Therefore, exacting comparative assessments of tools, like the SV mergers used in this study, are required.

By comparing SVCROWS with several modern and widely used SV merging algorithms, we showed that SVCROWS is as accurate as existing tools for validation (**Fig. 4**) while also being capable of maintaining diversity during SV discovery in a multi-sample dataset (**Fig. 3**). Additionally, optional functions of SVCROWS can be tuned to overcome unique challenges presented by datasets (**Fig. 2**) or provide different resolutions of complex regions in the dataset. Ultimately, it is the user’s choice as to whether it is more important to remove or maintain variation in the final SVR dataset. Because of this choice and its potential impact, no merger can be said to be objectively ‘better’. Instead, we emphasize the importance of options and balanced outcomes, which are the core principles of SVCROWS.

While some SV mergers are a part of more comprehensive pipelines, their merging function cannot be used independently of the rest of the pipeline, and/or the merging function has only been validated with defined upstream components(15–17). SVCROWS, on the other hand, is an accessible, standalone merger that is applicable to large or small datasets for both SV validation and discovery. To our knowledge, it is the only R-based SV merger, requiring minimal computational expertise. Further, the text-based input format of SVCROWS is flexible enough to incorporate all data types and formats. Finally, SVCROWS construction around RO as the main means of determining SV matching (i.e. no assignment of arbitrary ‘scores’ or ‘weights’) provides a level of transparency as to what caused two SVs to match or not. Given its flexibility the program also has appeal for more general applications of comparing sets of any genomic intervals, not just SVs.

Overall, we demonstrated that there are disparities between SV merging algorithms. The choice of which to use is up to the researcher, but we present three considerations based on direct comparisons of SV merging performance on both simple and complex datasets (**Fig. 5**). The first consideration is that the output of the program must reduce the complexity of the region (**Fig. 5**, ‘Output’). In this analysis, all mergers were able to reduce SV redundancy; however, most failed to strike a balance between under- or overmerging especially when a range of SV sizes were found in the same genomic neighborhood (e.g. Jasmine, Truvari, and Survivor, **Fig. 2C** and **4D**). We demonstrated above that this is less problematic for SVCROWS (**Figs. 2K, 3E** and **S5F**). A second consideration is that the architecture of the algorithm needs to be compatible with the study design (**Fig. 5**, ‘Technical’). The user should recognize that not all mergers work on all platforms, nor are they compatible with all upstream steps. A third consideration is that the algorithm type determines how the input SVs contribute to the resulting SVR (**Fig. 5**, ‘Algorithm Type’). For example, Jasmine includes RO as an optional parameter but prioritizes size and start location for merging decisions. Conversely, SVCROWS prioritizes RO and better reflects the relationship between SV size and potential functionality of those SV.

**Figure 5.**
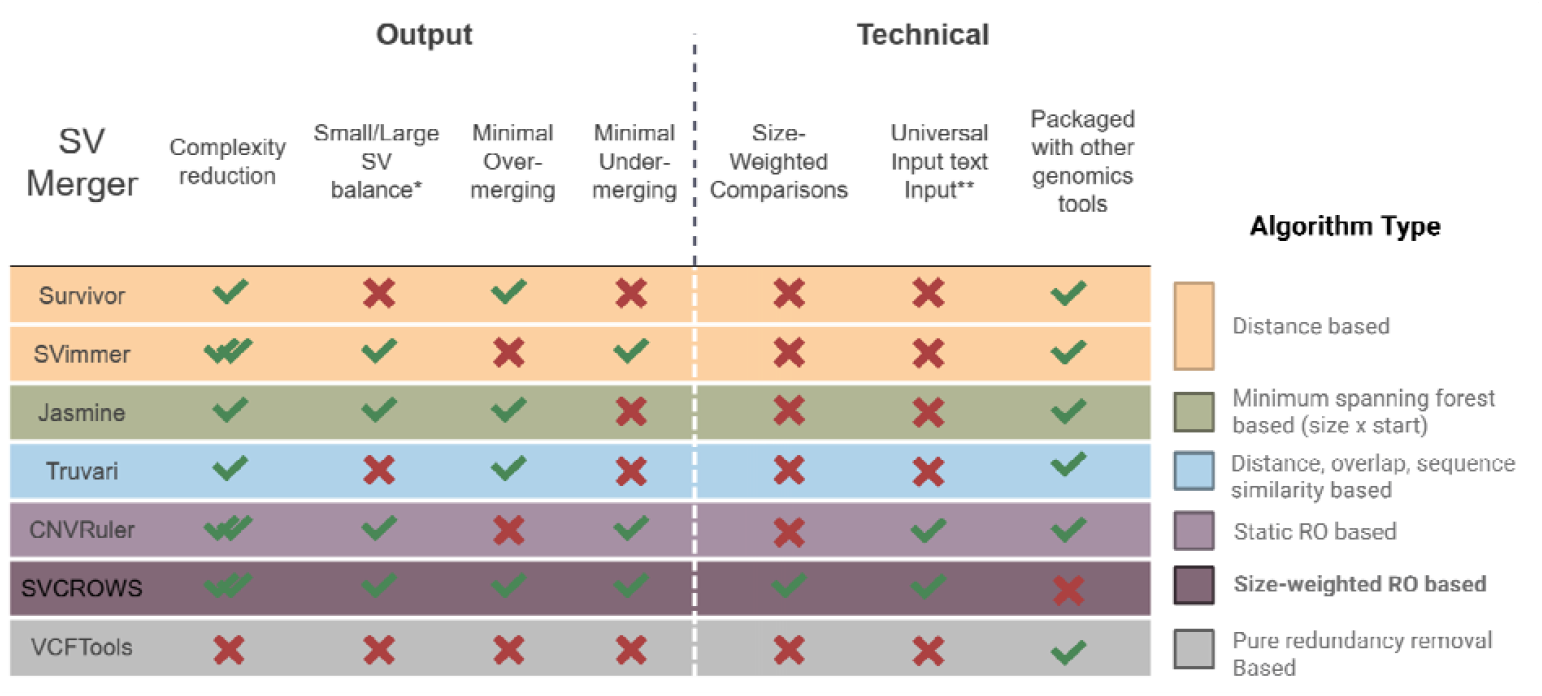
Comparison of tools based on proposed considerations to make when choosing an SV merging algorithm. SV merging evaluation of different features of each program (left); Key for algorithm each type of program uses (right).

SVCROWS was recently employed for SV discovery on a highly variable low-input genomics dataset and facilitated novel biological conclusions(54). Here, we also showed how merger choice can affect data reporting under biological contexts (**Fig. S8, 3A, 4C**), but correct merging can also aid in supporting causal relationships between genomic features. For example, a particular region on chromosome 16 in the ovarian/breast cancer SC dataset contained many SVs (**Fig. S11**). Upon further investigation, this region was part of a known super cis-regulatory elements (SCRE). SVs in SCREs can rapidly alter expression patterns in ways that drive developmental changes, phenotypic diversity, and are associated with worse cancer outcomes(55). According to the ENCODE database(56, 57), these SVs in the SC dataset fell into regions responsible for both upregulating and downregulating 4 genes immediately downstream(57). All 4 of these genes are implicated in ovarian and breast cancer in disparate capacities: some are more highly transcribed in ovarian or breast cancers, while others are less transcribed (**Fig. S11**)(58–61). Importantly, by preserving the variation across this region, SVCROWS helps convey that SVs called within the SCRE experience populational heterogeneity and may contribute to oncogenicity through diverse mechanisms, which in turn, can inform future experimental studies.

We do recognize that there are some limitations to SVCROWS implementation. While the SVCROWS’ input format is flexible, manual conversion from other formats may be necessary and add extra steps to a pipeline. Fortunately, SVCROWS includes quick-conversion tools to go between SVCROWS-input, BED, and VCF formats. Additionally, because SVCROWS makes more comparisons-per-SV to ensure accuracy, its computational time is longer than that of similar programs (**Fig. S3**). While running SVCROWS in R is more accessible, R-based software is notably slower than other languages. Further, the SVCROWS algorithm does not include sequence information during merging decisions. Other tools like Truvari or PanPop(16) can evaluate sequence identity at single base pair resolution in certain SVs (e.g. >50bp insertions and deletions), which may be more beneficial for studies where variance is low across large populations. However, these mergers are not compatible with all upstream calling tools required to achieve that level of precision. Further, a granular analysis of their ability to resolve complex regions (such as in **Fig. 3**) was not provided in those previous studies.

While these SV merging tools have been used to great effect in the past, SVCROWS promises improvements to the field. This is especially true given the growing number of large, multi-sample datasets that are harder to computationally refine accurately. Similarly, having tools like SVCROWS that are capable of resolving uncommon mutations in complex genomic regions will be of continued importance to researchers in need of high precision tools.

## Data Availability

SVCROWS can be downloaded as a Package or R-Markdown file at: https://github.com/A-Crow-Nowhere/SVCROWS.git

The source code for SVCROWS can be accessed at https://doi.org/10.6084/m9.figshare.29099801.v1

Links to the GIAB project and EGA data files can be found in the *Supplemental Methods*.

## Author Contributions Statement

Noah Brown: Conceptualization, Data curation, Investigation, Formal analysis, Software, Validation, Visualization, Writing-original draft, writing-review and editing

Charles Danis: Data curation, Formal analysis, Software

Vazira Ahmedjanova: Software

Jennifer Guler: Conceptualization, Funding acquisition, Methodology, Project administration, Resources, writing-review and editing

