## Supplementary Figures for "SVCROWS: A User-Defined Tool for Interpreting Significant Structural Variants in Heterogeneous Datasets"

**Supplemental Figures**

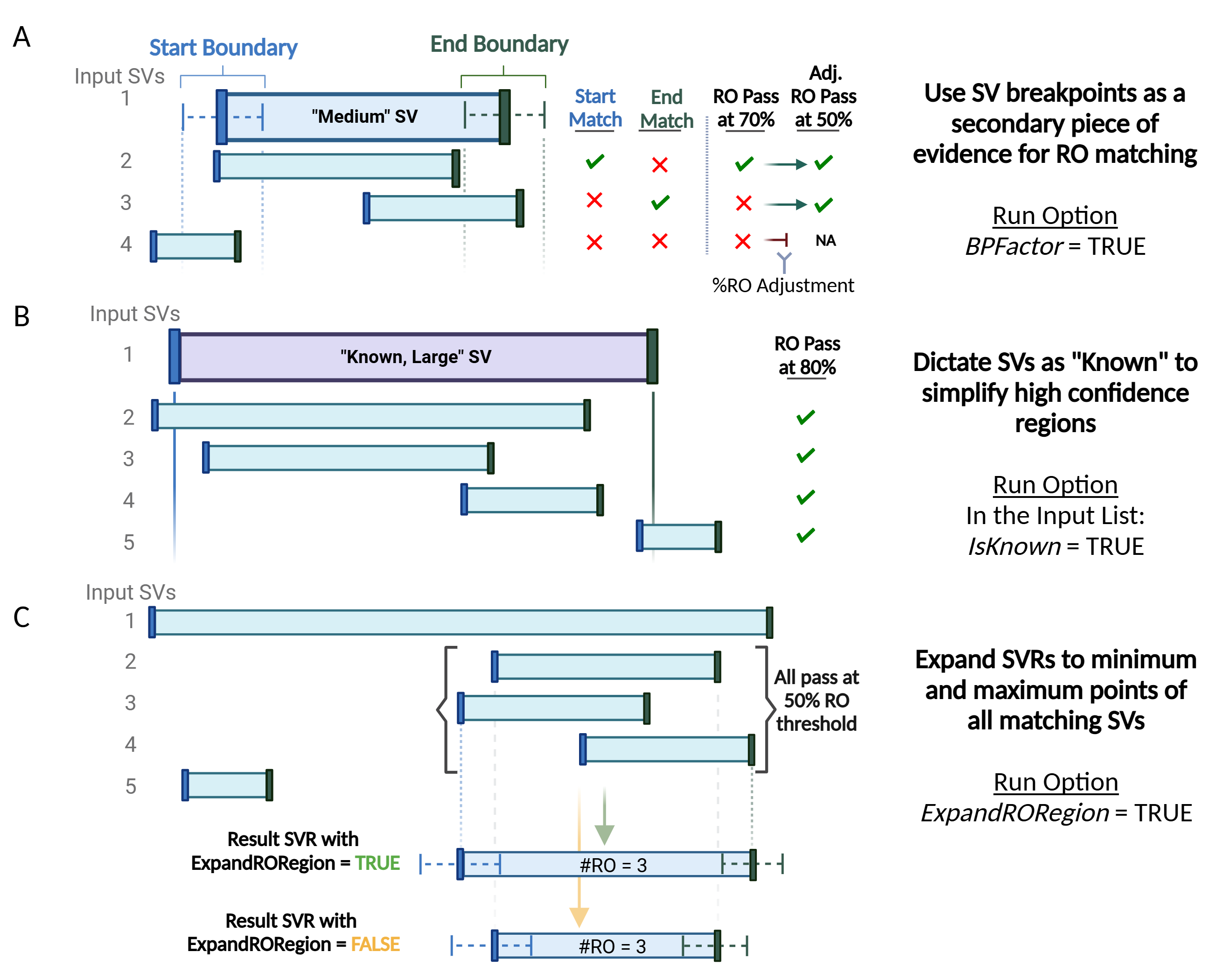

**Supplemental Figure 1. Optional calculation methods of SVRs within SVCROWS.** Any combination of these options can be applied. **A)** Breakpoint matching should be included as secondary information for SVR calling. With this option enabled, when subsequent SVs are compared, they check both the ‘Start’ and ‘End’ breakpoints for overlap with the region defined by the user (Inputs 3&4). If a match is found (green check), the resulting %RO threshold is decreased to the minimum used in the analysis (in this case, 50%). **B)** Designation of input SVs as “Known”. When SVs are marked as known, any subsequent SV compared to it will have an overlap requirement of 1 base pair. “Known” SVs are queried against first in the analysis, so no new SVRs are built in those regions. **C)** Expansion of SVRs to maximum end points. By default, SVCROWS will use the boundaries of the largest SV in an SVR to define the region, With the ExpandRORegion option enabled, it will instead expand the region to the most distal breakpoints of that region and update the breakpoints.

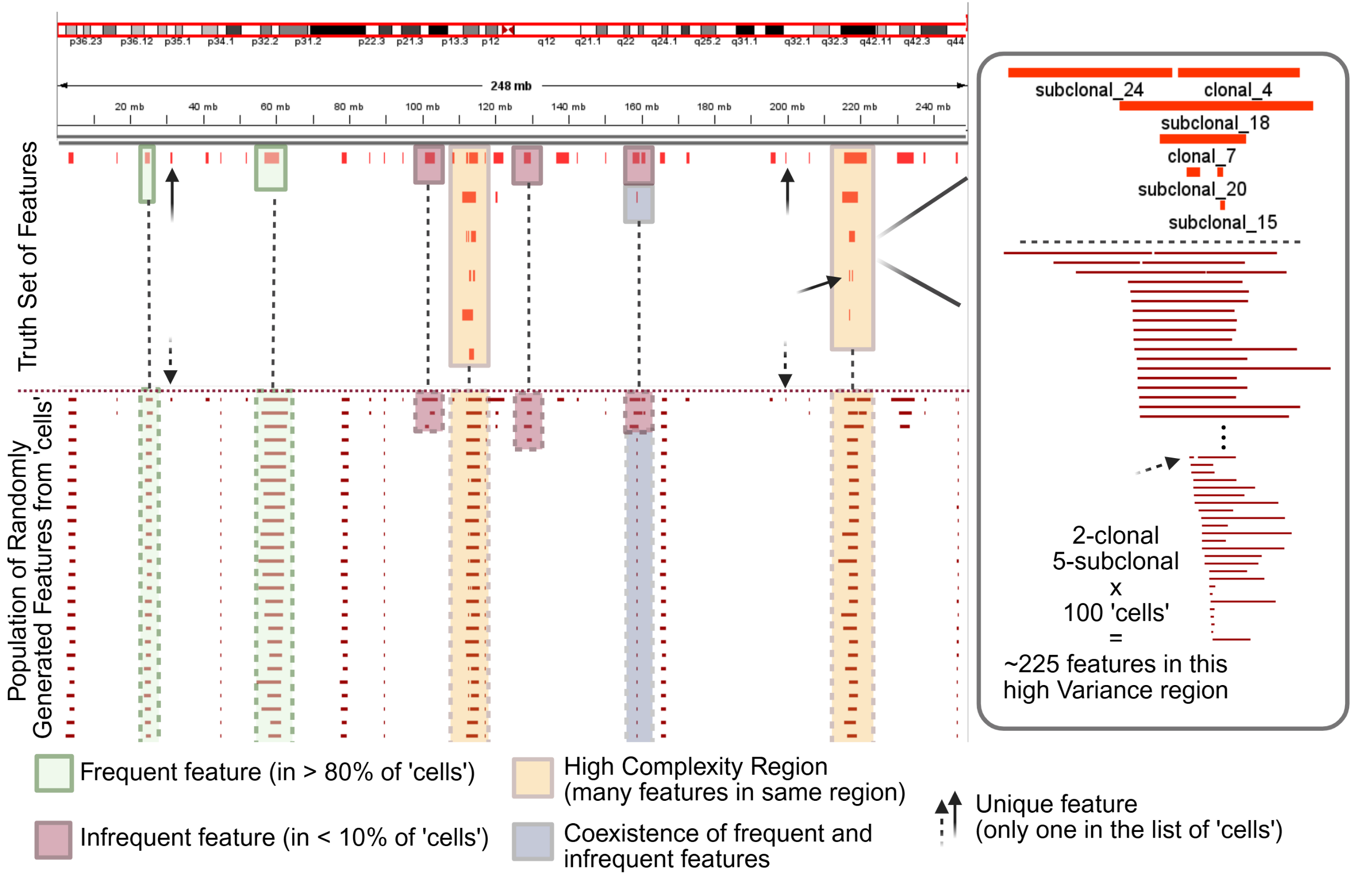

**Supplemental Figure 2. Annotated IGV view of a SV-Simulation run.** (**Left** top): the full human chromosome 1 (hg38) with 50 randomly generated total Truth Set features that mimic SVs in potential datasets. Each features has a known frequency across the entire simulation prior to ‘cell’ generation (**Left,** bottom): a concatonated list of the simulated ‘cells’ that are made to reflect the pre-determined frequencies of the Truth Set (note variance in breakpoint position). Examples of different features in the SV-Simulation are represented with by Truth Set Feature(s) (solid box) and an experiment wide list of the simulated ‘cell’ feature(s) (dashed box): Clonal (green) - features that are present commonly in the population of ‘cells’, representing germline SVs. Subclonal (red) - features that occur rarely in the ‘cell’ population, representing easy to miss rare SVs that are sometimes merged out of the data. Clonal and subclonal Truth Set features can randomly overlap (blue), leading to mixes challenging to reslove correctly with an SV merger. The SV-Simulation also specifies wither to inlude high complexity regions (yellow), which intentionally overlaps a specified number of features. Arrows denote SVs that occur only once in the simulated ‘cell’ population. Solid arrows point to the correct, Truth Set position while dashed arrows point to the corresponding existance in the population. (**Right**): a detailed view of one of the high complexity regions.

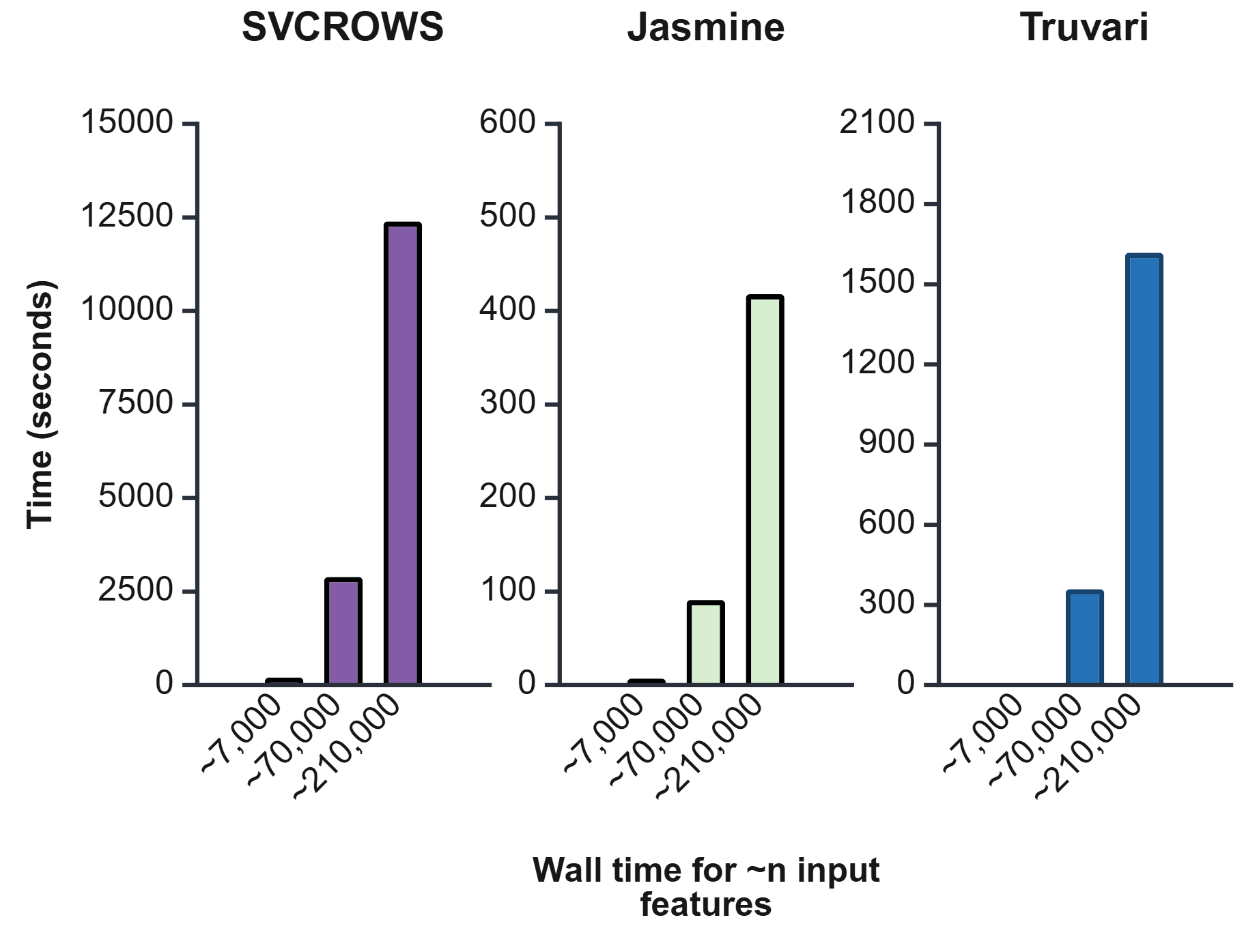

**Supplemental Figure 3. Comparison of SV-Merging programs over n number of features.** Inputs for these tasks were constructed from SV-Simulation. N=1 for all time points. Processor: Intel(R) Core(TM) i7-8750H CPU @ 2.20GHz RAM: 16.0 GB. System Type: Windows, 64-bit operating system, x64-based processor.

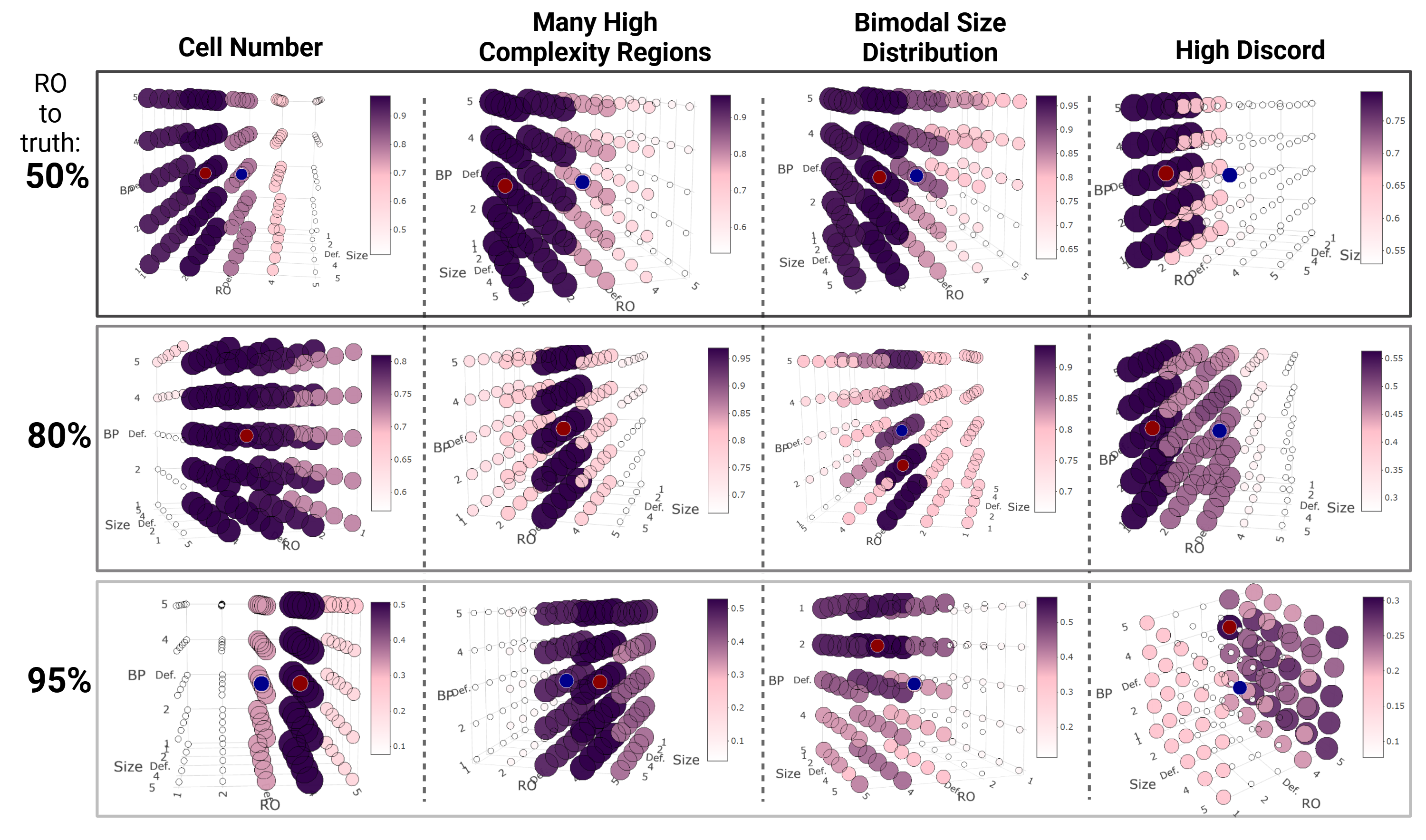
 **Supplemental Figure 4. 3D visualization of F1 scores of SVCROWS-Default over multiple challenging SV-Simulation datasets.** Highlighted blue dot represents the default F1 score, while the red dot indicates the highest F1 score after parameter optimization. Full input specficiations for all runs, and full results for the 95% RO can be found in **Table S1.**

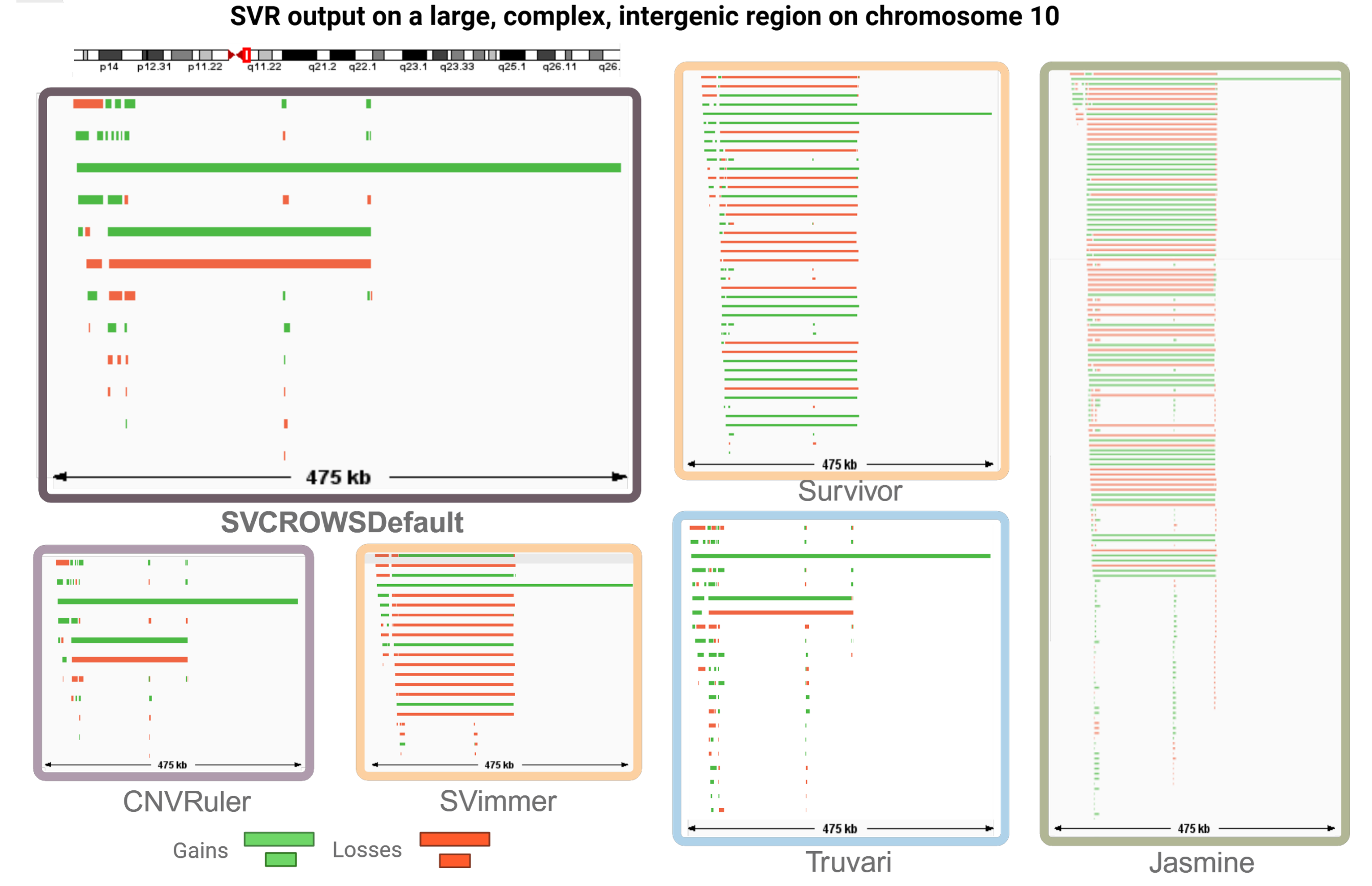

**Supplemental Figure 5. Region specific ccomparison of SVCROWS to 5 SV merging programs using a heterogeneous SC dataset.** Integrated Genome Viewer visualization of resulting SVRs in the chr10 region for each program.

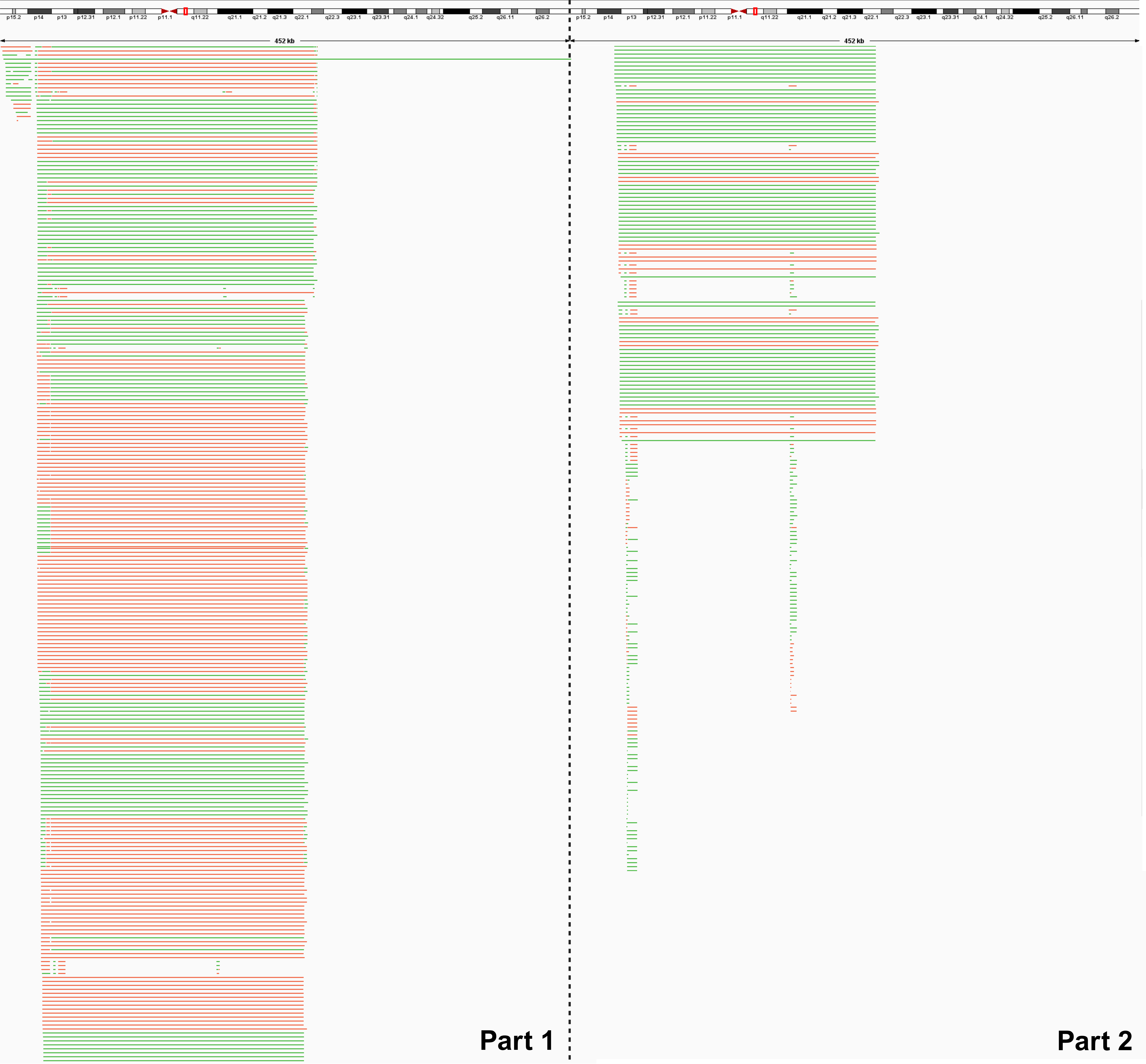
**Supplemental Figure 6. IGV view of the VCFTools merging of the complex chromosome 10 region from the ovarian cancer dataset.** Complex regions such as these are difficult to merge. Red feautres represents sequence loss SVR, while green indicate gain SVRs. N = 804 SVRs.

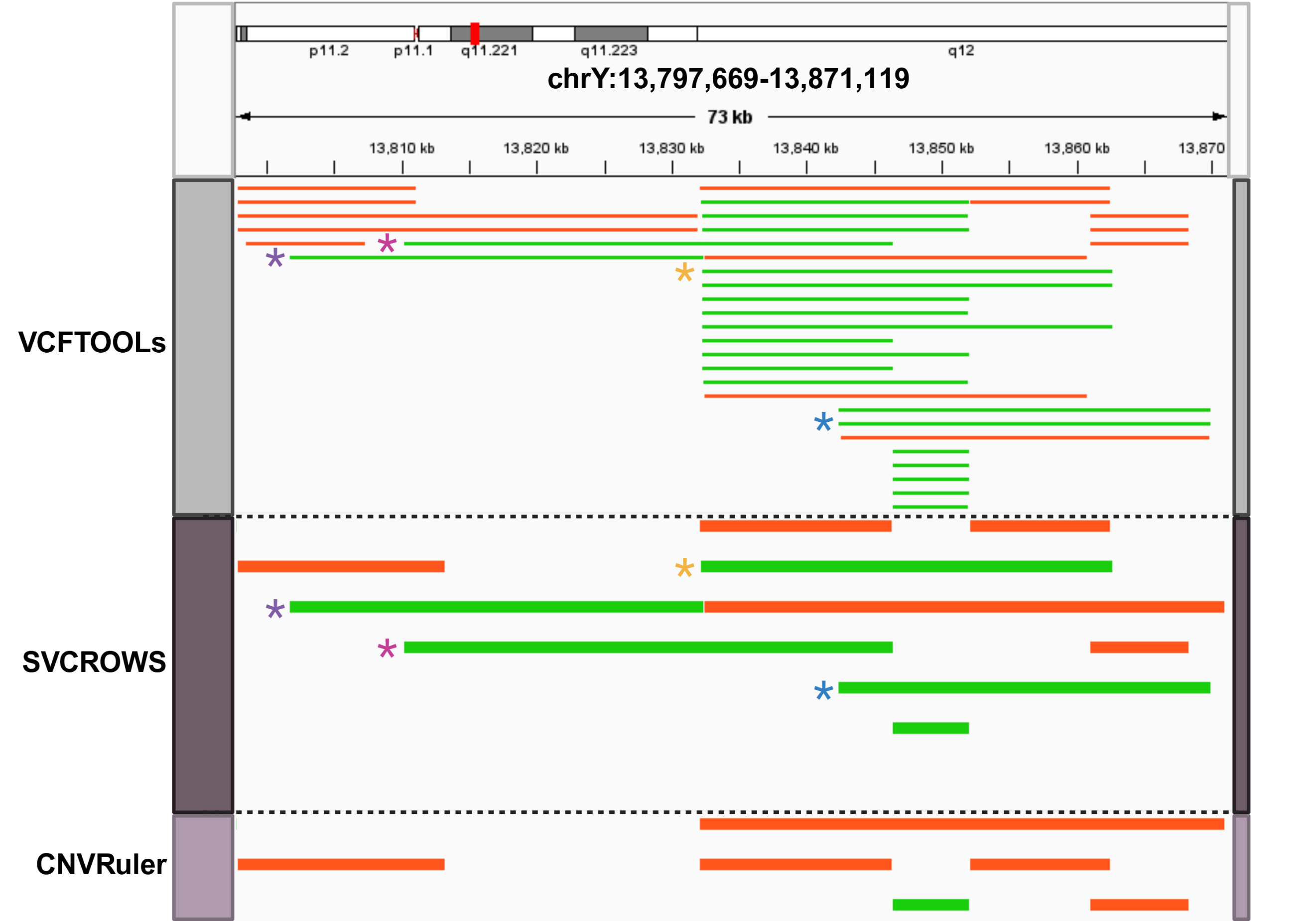

**Supplemental Figure 7. IGV view of a region with both smaller and larger SVRs in the HG002 benchmarking dataset.** Direct vizualization of these tools shows that CNVRuler missess several key SVRs (Dentoted by *) that are reported in VCFTools (Pure redundancy removal). For the purposes of vizualization, SVRs >40kb were hidden from the viewing pane.

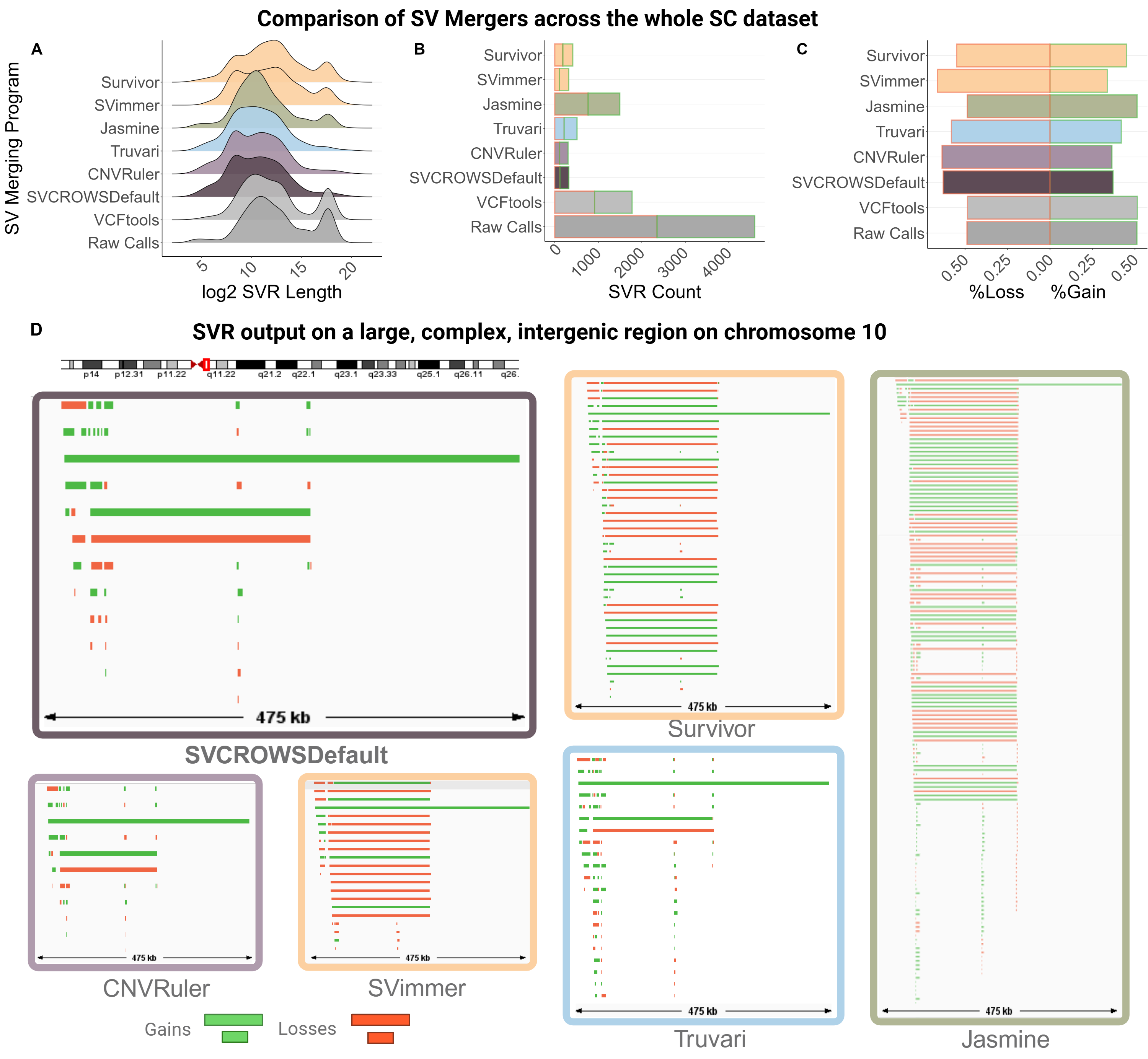

**Supplemental Figure 8. Dataset-wide comparison of SVCROWs to 5 SV merging programs using a heterogeneous SC dataset.** SV merging performance of different algorithms using the single cell ovarian cancer dataset. **A)** Size distribution of resulting SVRs. **B)** Final SVR count of each tool, compared to the unmerged raw lumpy calls. **C)** Distribution of gains and losses in final SVRs.

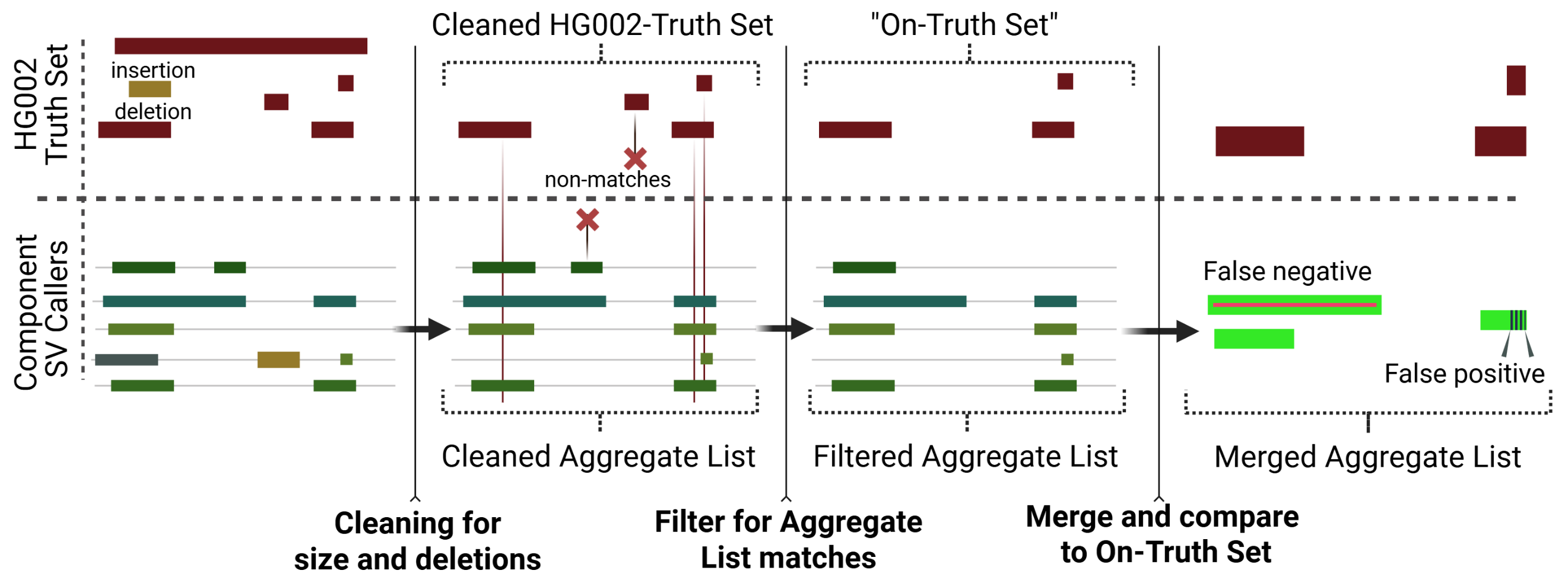

**Supplemental Figure 9. Diagram depicting the steps used to filter and implement the HG002-Truth Set for subsequent comparative analysis.** The three step process for making the On-Truth Set, which only contains SVs from the HG002-Truth Set (top, red) that are relevant to compare with those in the compliation of SV calls from 5 SV callers, termed the Aggregate List (bottom, green). The original SV calls from the GIAB consortium contain both deletions and insertions depending on the caller used. The Aggregate List contains some of the component SV calls that were used to make the HG002-Truthset, and some that are low accuracy calls (1^st^ panel). Step 1: for simplicty, SVs greater than 2Mb and insertions were excluded from the Aggregate list, and HG002-Truth Set. The result is the Cleaned HG002-Truth Set and Aggregate list; used in main-text panels **Fig. 4A-C** and **S10-11** (2^nd^ panel)**.** Step 2: Further filter the HG002-Truth Set and Aggregate List to create a the On-Truth Set (3^rd^ panel). SVs must have atleast 1 BP overlapping between the Aggregate List and Filtered HG002-Truth Set to be kept (panel 2, red X). Step 3: Merge the Aggregate List, and compare to the On-Truth Set; used in main-text panel **Fig. 4D** (4^th^ panel). The Merged Aggregate List may contain false negative or false positive merging calls depending on the merger and parameters used, which are factored in to the F1 score.

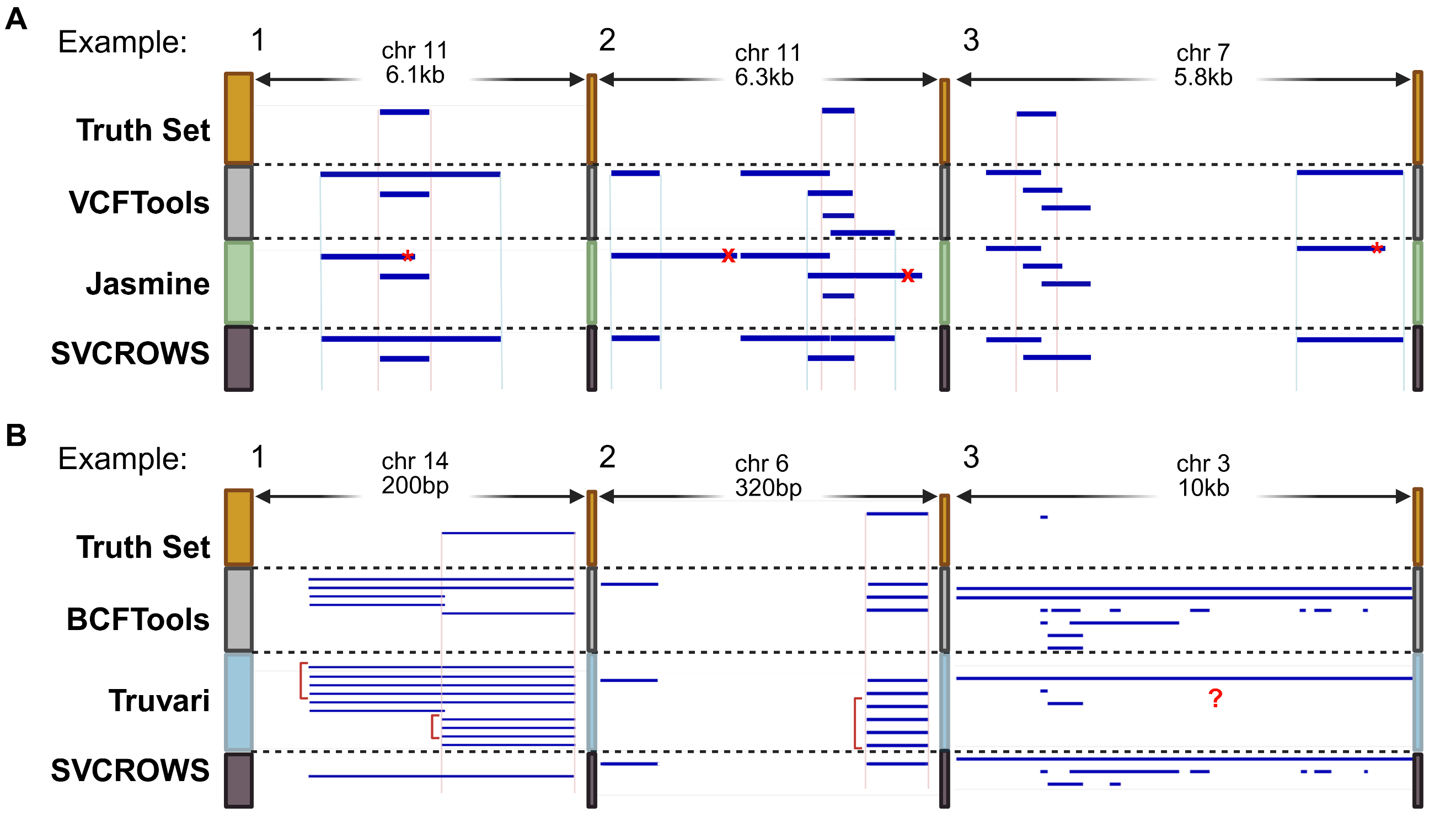

**Supplemental Figure 10. IGV view of comparing Jasmine and Truvari to SVCROWS in the resulting HG002 SVRs.** 3 separate examples are depected, all blue bars are deletions. The Truth Set (yellow) represents SVRs that have been validated by the Genome in a Bottle (GIAB) project through multiple callers. VCFTools or BCFTools represent the complete set of calls from the 5 chosen GIAB callsets, including those that did not pass quality thresholds to be included in the Truth Set. **A)** Comparison of Jasmine to SVCROWS. Pink lines represent the Truth Set breakpoints, while blue lines represent the correct non-truth breakpoints of included SVs, which can either be premature stops (red *), unexpected elongations (red ‘x’). Example 1; Both SVCROWS and Jasmine match the Truth set, but Jasmine has a premature stop in the non-truth merge. Example 2; Jasmine maintains closer similarity to the truth set, but experiences two unexpted elongations in the complex region. Example 3; Jasmine and SVCROWS both are innacurately merged relative to the Truth Set, but Jasmine experiences a premature stop in a unrelated location. **B)**  Comparison of Truvari to SVCROWS. BCFTools is used instead of VCFTools, at is is the reccomended option for Truvari. Pink lines represent the Truth Set breakpoints, while red brackets represent artifactual SVRs. Example 1; Truvari is more acruate to the Truth Set, but creates artifactual SVRs. Example 2; Truvari and SVCROWS both match the Truth Set SVR, but Truvari introduces more artifactual SVRs. Example 3; Truvari loses SVR heterogeneity found in the input across a complex region.

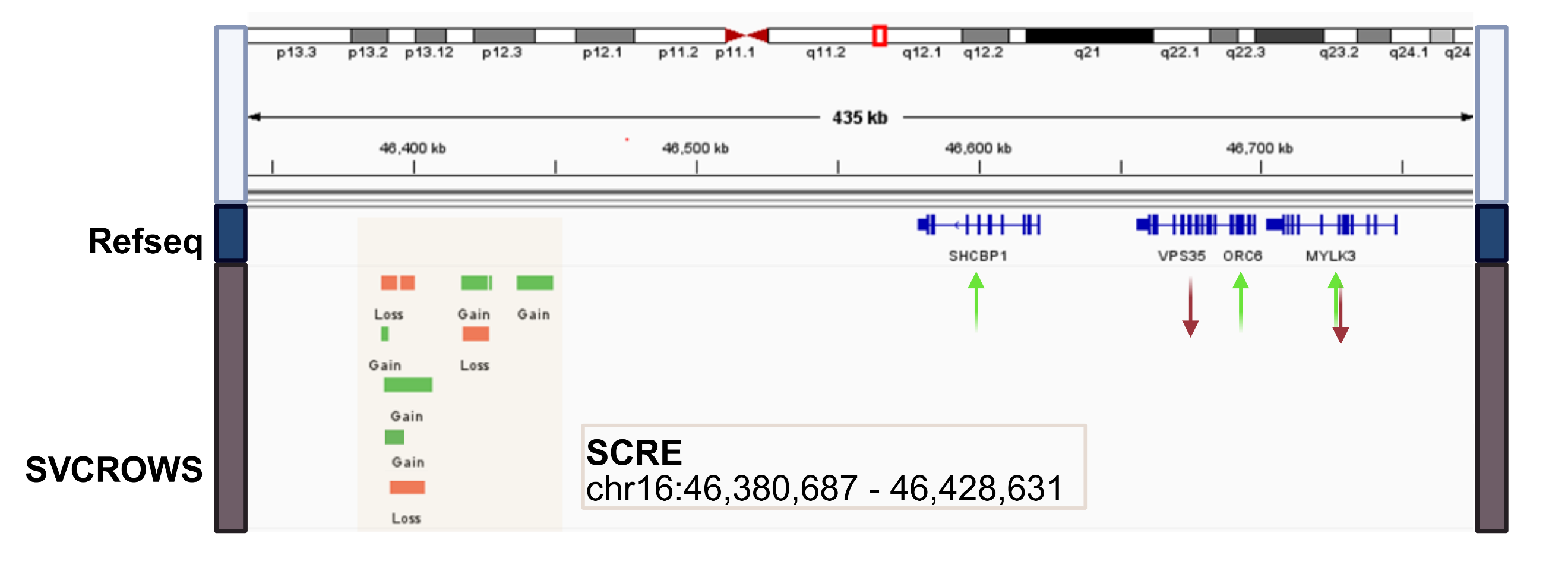

**Supplemental Figure 11. IGV view of a complex region merged by SVCROWS and the downstream genes.** A portion of chromosome 16 from the SC ovarian cancer dataset. The SVRs fall into a known Super Cis-Regulatory Element (SCRE) region which has been documented to have direct impacts in transcriptional output of the genes directly downstream. Some genes have been documented to be tumor supportive (green arrows), while others are tumor repressive (red arrows) when over transcribed in several forms of cancer including ovarian cancer and triple-negative breast cancer. SVCROWS’ ability to preserve heterogeneity while reducing complexity demonstrates how a SCRE may be heavily selected in individual cells for certain genotypes.
