## Supplementary Methods for "SVCROWS: A User-Defined Tool for Interpreting Significant Structural Variants in Heterogeneous Datasets"

**Supplemental Methods**

**Detailed SVCROWS components**

**Usage**

SVCROWS can be downloaded as an R package from https://github.com/A-Crow-Nowhere/SVCROWS.git All functions in the application require an input list (in a .bed/.tsv format) to be provided with the parameters in **Table M1**. For the analysis, the user combines all SVs detected in the dataset (e.g. in a single cell analysis, every cell and every SV called). Each entry in the input must contain all of the following information. All String type variables have no ‘fuzzy’ matching tolerance (i.e. “Chr1” ≠ “chr1” and “Deletion” ≠ “Del”).

SVCROWS will loop through an entire directory of input files for processing. The application will stop if a non-Input List format file is present in the input directory. Output files are written based on the first 15 characters of the input file and renamed into their respective file types.

**Table M1: Input table format**. Blue are features typically calculated from other SV-calling software. Green are run specific information provided by the user. Yellow are optional calculations that are made alongside each run.

| **Column Header** | **Type** | **Description** |
| --- | --- | --- |
| *Chr* | String | Chromosome of SV |
| *Start* | Int | *Start* (5’) of SV *Cannot be negative |
| *End* | Int | *End* (3’) of SV *Cannot be negative |
| *Length* | Int | *Length* of SV used to place in size categories and calculate RO *Can be negative (I.e. for a genetic deletion) **Will mask *End*-*Start* from above (i.e. must match the length given from *Start* and *End*) |
| *Type* | String | The *Type* of genomic event occurring in this region (e.g. del, dup, inv) *Multiple *Types* can be assessed simultaneously, but will not match each other if RO regions overlap. |
| *ID* | Int | Numeric *ID* given to each entry in the Input list *Must have no repeat IDs |
| *Var1* | String/Int | Can contain whatever relevant information for downstream analysis, and is not used in any calculation. |
| *Var2* | String/Int | Can contain whatever relevant information for downstream analysis, and is not used in any calculation. Include VCF;INFO field information here. |
| *Var3* | String/Int | *Var3* can also be considered a “sample_ID” for the dataset. Use this column to denote different individual samples (not different SVs) in the data. The program will eventually use this variable to calculate each sample’s contribution to the final SVRs. |
| *IsKnown* | Boolean | Is Known will either be “TRUE” or “FALSE” for each sample, see the feature description for its function. For all default functionality, use “FALSE”. |
| *NumReads* | Double | This column intends to summate the number of reads used to call each SV matching to an SVR, but a user can input any relevant information to add together. If the user has nothing to compute here, they can input “0”. |
| *QScore* | Double | This column intends to average the quality score associated with each SV matching to an SVR, but a user can input any relevant information to average together. If the user has nothing to compute here, they can input “0”. |

If a population-level analysis is required, it may be beneficial to pre-merge features within samples. For example, a SV may be called twice with slightly different breakpoints within the same sample (reflecting the same biology, but present only due to technical error). So that they are not ‘double counted’ in a populational merge, the user can:

1. prepare all individual samples as SVCROWS input format (easily converted from BED and VCF using included tools)
2. run a within-sample merge for each sample in the input directory (need only run once)
3. Concatenate output FCL (see below) files into a single ‘cleaned’ list
   a. Optionally ensure each SV from each sample has a key for recognizing that sample in the Var1,2,3 columns, so sample fates can be tracked
   b. FCLs formatted output files (see below) can be used as input for this reason
4. run SVCROWS again on the single, between-sample file
5. the resulting files are then representing cleaned, population-level data

The application can be run on the R console as such:

Scavenge(InputQueryList = "~/user/R/SVCROWSin", OutputDirectory = "~/user/R/SVCROWSout", ExpandRORegion = FALSE, BPfactor = TRUE, DefaultSizes = FALSE, xs = 5000, xl = 25000, y1s = 500, y1l = 2500, y2s = 50, y2l = 80)

**User Parameters**

Along with the input file containing potential SVs, the user must also provide 6 parameters to generate the SVCROWS algorithm, which will set the stringency for matching in SVs of varying sizes (**Fig. 1C** purple line). Stringency is user-defined for each parameter. The line generates a dynamic set of values used for SV overlap calculation, using *y = mx + b,* where *x* is the length of the SV, and *y* is the resultant stringency value. Two different y-axis are used, but follow the same pattern based on the shared x-axis.

i) Firstly, the user must define limits on size categories for the SVs to create ‘small’, ‘medium’, and ‘large’ (**Fig. 1C,** inputs 1&2). The user makes the choice based on the distribution of the sizes of the SVs in the input file, which is typically positively skewed (**Fig. 1C,** teal bell curve). ‘Small’ size SVs consider any variation as non-biologically impactful, and therefore SVs in this category have a minimum and constant set of parameters. Conversely, because matching ‘large’ SVs have such high levels of biological similarity, variation is largely considered to be caused by SV calling, not true biological differences. Therefore, further size increases do not affect the resulting variables. As a consequence ‘medium’ SVs have varying stringencies of other variables based on their length.
ii) Secondly, the user provides the length of a boundary, centered around either breakpoint at the ends of an SV, that defines a region where potential SV’s breakpoints will be considered matching (see **Fig. S1A**). The user defines these boundary sizes at the ‘small’ and ‘large’ categories, which then sets the size range to be calculated for the ‘medium’ category (**Fig. 1C,** left y-axis and inputs 3&4).
iii) Finally, a user provides a RO threshold at ‘small’ and ‘large’ size categories limits as a percentage. This most directly controls the stringency of matching SVs (**Fig. 1C,** right y-axis and inputs 5&6)

**Detailed feature descriptions**

i) SVCROWS can generate default values for these 6 parameters based on the second and fourth quartile of the SV-size distribution of the dataset. It is important to note that using this option will adjust its parameters to each new file in the input directory. However, use of this option should be taken with caution, as it may not tailor the application to user needs.
ii) SVCROWS uses breakpoint matching between SVs as a secondary piece of information to determine the similarity between two SVs (**Fig. S1A**). The user input determines breakpoint boundary sizes. As SVs are compared, the start or end breakpoint must fall within the boundary to be considered a match. Start boundaries cannot match with end boundaries and vice versa. When a breakpoint matches, SVCROWS can interpret this as a high likelihood of a true biologically similar SV and adjusts the RO threshold to the minimum value (**Fig. 1C,** input 5) provided by the user (**Fig. S1A,** green arrows). If potential SVs are already ‘small’, there is no change. Breakpoint matching is counted and recorded on the consensus list only once to the first match found in the consensus list (so, if multiple breakpoints align, they will all go to the same resultant SVR), but subsequent matches will continue to adjust RO thresholds after it is counted.
iii) The input SV list required by SVCROWS has an optional distinction to indicate a potential SV as a “Known” SV. This characterization reduces the RO requirement to a single base pair (**Fig. S1B**). For example, this option can be used for SVs that have been previously identified and have definite positions in the genetic background of an organism. Therefore, a user can have high confidence that any matching SVs in this region are derived from the “known” SV, and any variation in SVs compared in this region is not biological – or remove any variation in a region.
iv) In many analyses, the user wants to first merge a host of SVs from different samples, and then keep track of the relative contribution/outcome of SVs per sample. SVCROWS provides an output file that mirrors the input list, but includes the merging fate, and its relative weight in the SVR outpout. See the description of the PerSampleList and AdjustedPerSampleList below
v) SVCROWS provides an option to expand SVRs with multiple matches to the minimum and maximum position of the subset of matching SVs (**Fig. S1C**). Using this option adjusts the breakpoint regions for the newly constructed region during the run itself. Breakpoint matching is still tabulated in the same manner with the adjusted values.
vi) SVCROWS includes an alternative function (“Hunt”), where a user inputs both 1) a set of SVs or SVRs in the same input format as described for “Scavenge” mode, and 2) A set of features (i.e. genes from a relevant genome). Using the same weighted-sizes principle, SVCROWS will quantify how many times features are significantly overlapped by the supplied user input SVs. However, this calculation is done non-reciprocally and only considers the overlap of the features to call matching SVs. The logic of the comparison to match is:

(#Of_Feature_BP / #Of_SV_BP) > RO-Threshold

The ‘feature list’ has the same requirements as the ‘input list’ but only contains the: *“Chr”, “Start”, “End”, “Type”, “Var1”, “Var2”, and “Var3”* headers (see https://github.com/A-Crow-Nowhere/SVCROWS.git for an example).

During runtime, the user must provide the same 6 variables as in “scavenge” mode, but consider inputs that apply to the characteristics of the features themselves, rather than the SVs. This mode returns similar outputs as the “Scavenge” mode, but the consensus list represents the quantification of each feature given to the application.

vii) There are several tools and functions for quantifying the resulting SVRs. Incorporated into the ‘input list’ are the variables *“NumReads”* and *“QScore”*, which are added and averaged (respectively) as individual SVs match into their final SVR (see **Usage** for further functionality). SVCROWS also has a “Summary” function, that will provide summary information of the final SVR output.

**Output Files**

SVCROWS has four output files with various information.

i) QueryList: An ordered version of the input list (**Table M1**). Reflects the order the SVs are compared. (File extension = .QL)
ii) ConsensusList (**Table M2**): The principle output of SVCROWS; it shows unique SVRs extracted from the input list. The number of matches to the SV in the consensus list across the input dataset is reflected by the ‘ROPass’ column. The frequency and rarity are calculated and assigned based on the number of unique samples in the input dataset (*Frequency* = #ofmatches/#uniquesamples). (File extension = .FCL)
iii) PerSampleList (**Table M3**): SVCROWS tracks the outcome of each input SV. It is constructed in the same as the QueryList with 4 extra columns for assigning the ID number of the ConsensusList entry it matched with, or its own ID number if it was unmatched. Breakpoint matching is carried out similarly. Rarities are determined based on ID matching from the ConsensusList. (File extension = .FPSL)
iv) AdjustedPerSampleList: This output is a version of the PerSampleList that removes duplicate matches to the same ConsensusList entry within a sample. For example, if sample-A has two SVs called in a region (likely reflecting the same biology), that both match to SVR1, only the larger will be indicated as matching to SVR1. The other SV will be marked as ‘clone’, and have a negative matching ID number (essentially, to be excluded from downstream analysis). Based on user preference, this may reflect more closely the true nature of SVs in a dataset. (File extension = .AFPSL)

**Table M2: FinalConsensusList Format**. Gray rows follow the same description as in the input list (**Table M1**) unless otherwise indicated. Orange rows are calculated by the program as a means of describing the SV. Green rows are quantitative results matching. Blue rows are post-hoc additions to further analyze the data.

| **Column Header** | **Type** | **Description** |
| --- | --- | --- |
| *Chr* | String |  |
| *Start* | Int |  |
| *End* | Int |  |
| *Length* | Int | *Length* of final SVR |
| *Type* | String |  |
| *ID* | Int | Final SVR will use the *ID* of the largest SV in the SVR |
| *Var1* | String/Int | Final SVR will use the *Var1* of the largest SV in the SVR |
| *Var2* | String/Int | Final SVR will use the *Var2* of the largest SV in the SVR |
| *Var3* | String/Int | Final SVR will use the *Var3* of the largest SV in the SVR |
| *IsKnown* | Boolean |  |
| *NumReads* | Double | Final Sum of all matching SVs in the SVR |
| *QScore* | Double | Final Average of all matching SVs in the SVR |
| *Size* | String | Which size category the SVR falls under |
| *BPBoundSize* | Int | Total length of the Breakpoint matching region (set by the user) |
| *ROPercentPass* | Int | The RO threshold needed for two SVs to match at this SV length |
| *ROCount* | Int | Tabulation of each matching SV in the SVR |
| *BPStartLeft* | Int | The left-most boundary of the breakpoint on the 5’ end of the SVR (Start) |
| *BPStartRight* | Int | The right-most boundary of the breakpoint on the 5’ end of the SVR (Start) |
| *BPStartCount* | Int | Tabulation of each matching *Start* boundary across all SVs in the dataset |
| *BPEndLeft* | Int | The left-most boundary of the breakpoint on the 3’ end of the SVR (End) |
| *BPEndRight* | Int | The right-most boundary of the breakpoint on the 3’ end of the SVR (End) |
| *BPEndCount* | Int | Tabulation of each matching *End* boundary across all SVs in the dataset |
| *Frequency* | Double | How many invidual SVs matched to this region.  Calculated as: *ROCount/(total unique values in Var3 column)*  *To avoid double counting in a region, it is suggested to run each unique sample individually in SVCROWS to consolidate complex regions |
| *Rarity* | String | Categorically assigned: *Frequency = 1/totalUniqueVar3, Rarity = ‘Unique’;*  *Frequency <10%, Rarity = ‘Rare’; Frequency >= 10%, Rarity = ‘Common’; If IsKnown = True, Rarity = ‘Known’* |
| *Matches* | String | Concatenates IDs as SVs match into a single string with a ‘:’ delineator. If *ExpandRORegions* = TRUE, if breakpoints are changed the delineator becomes a ‘+’. |

//

**Table M3: FinalPerSampleList Format**. Gray rows follow the same description as in the input list (**Table M1**) unless otherwise indicated. Orange rows are ParentIDs of the resulting matches from the FinalConsenssusList. Green row is based on the output of the FinalConsensusList.

| **Column Header** | **Type** | **Description** |
| --- | --- | --- |
| *Chr* | String |  |
| *Start* | Int |  |
| *End* | Int |  |
| *Length* | Int |  |
| *Type* | String |  |
| *ID* | Int | *IDs will be reordered in decreasing *Length* |
| *Var1* | String/Int |  |
| *Var2* | String/Int |  |
| *Var3* | String/Int |  |
| *IsKnown* | Boolean |  |
| *NumReads* | Double |  |
| *QScore* | Double |  |
| *BPStartParentID* | Int | The ID of the entry of the SVR Start-Breakpoint that this SV matched to. If it is the same ID as the ID contained in the PerSampleList, this indicates that this feature was unique. |
| *BPEndParentID* | Int | The ID of the entry of the SVR End-Breakpoint that this SV matched to. If it is the same ID as the ID contained in the PerSampleList, this indicates that this feature was unique. |
| *ROParentID* | Int | The ID of the entry of the SVR that this eventually SV matched to. If it is the same ID as the ID contained in the PerSampleList, this indicates that this feature was unique. |
| *Rarity* | String | The eventual categorization of the rarity of this input SV from the FinalConsensusList *in the AdjuststedFinalPerSampleList, If multiple SVs from the same sample (determined by values in *Var3*) match to an SVR, SVCROWS will only count display the matching rarity to the largest of those SVs. The others will be called “clone”. This mitigates potential issues in double counting. |

**Detailed description of SV-Simulation**

The code for this script can be found at <https://github.com/A-Crow-Nowhere/MalariAPI/tree/main/scripts/cnv_sim> . Example runs can be found in Table M4.

Canonical Event Generation

The simulator begins by generating a **canonical truth set** of CNV-like events, which represent the “ground truth” across the simulated cohort. Users specify the number of **clonal** (--n-clonal) and **subclonal** (--n-subclonal) canonical events.

- **Clonal events** receive prevalence values drawn uniformly from 0.80–0.99, ensuring that each event is present in the majority of cells.
- **Subclonal events** receive prevalence values derived from a geometric-like decay model (--p-geom), which favors rare events. Specifically, the simulator samples a cell-count *k* from a truncated geometric distribution, then sets the true frequency as *k / n_cells*. The parameter --max-subclonal-cells caps how widespread any single subclonal event can be.

Each event is assigned genomic coordinates on a chosen interval, a chromosome for example (--chrom), using lengths sampled log-uniformly between --min-size and --max-size. This distribution produces many shorter SVs with occasional large ones, consistent with real data. Overlaps among canonical events are permitted.

High-Complexity Regions

The simulator also creates high-complexity regions, which model loci prone to genomic instability. The user specifies the number of high-complexity regions (--hv-n-regions), their size (--hv-radius), and the minimum number of canonical events that must overlap at least one High Complexiy region (--hv-min-events). The simulator enforces this guarantee by repositioning selected canonical events so that they overlap the HV interval. It then optionally biases additional canonical events to overlap the other HV regions. Each canonical event records its membership in the output.

Per-Cell Assignment

For each canonical event, the simulator determines the number of cells it should appear in by rounding true_freq × n_cells. It then randomly selects that many cells without replacement. This procedure ensures that the per-cell counts reflect the target prevalence while introducing binomial sampling variation when viewed across multiple runs.

Discord Model

After assigning canonical events to cells, the simulator introduces positional and size jitter to mimic technical imprecision in breakpoint calling. A single run-level parameter, --variance (1–100), controls the magnitude of this jitter:

- 1 produces no deviation (observed = truth).
- 10–25 produces subtle shifts (often <1 kb for events in the 50–200 kb range).
- 40–60 produces moderate shifts (hundreds of bp to several kb) and 10–20% size changes.
- 70–100 produces large deviations (multi-kb shifts; 30–70% size changes).

Larger SVs experience proportionally greater absolute shifts. The jitter is applied symmetrically

(left or right shifts, lengthened or truncated), ensuring that not all errors skew in one direction.

Output Files

The simulator writes BED-formatted files to the specified output directory:

- canonical_events.bed — Truth set of CNVs with their true_freq, type (clonal/subclonal), and high-complexity region membership.
- per_cell_cnvs.bed — All observed CNVs across cells, with cell IDs.
- per_cell/*.bed — Per-cell CNV tracks.
- event_observed_freqs.bed — Table comparing expected vs observed event frequencies.
- high_variance_regions.bed — HV loci and the canonical events overlapping them.

All outputs are IGV-compatible and can be directly visualized.

**Example Runs of SV-Simulation**

**Table M4: Example command line runs for the simulated experiment replicates used in Figure 2.** Between replicates and experiments, the seed number and outdir are changed manually.

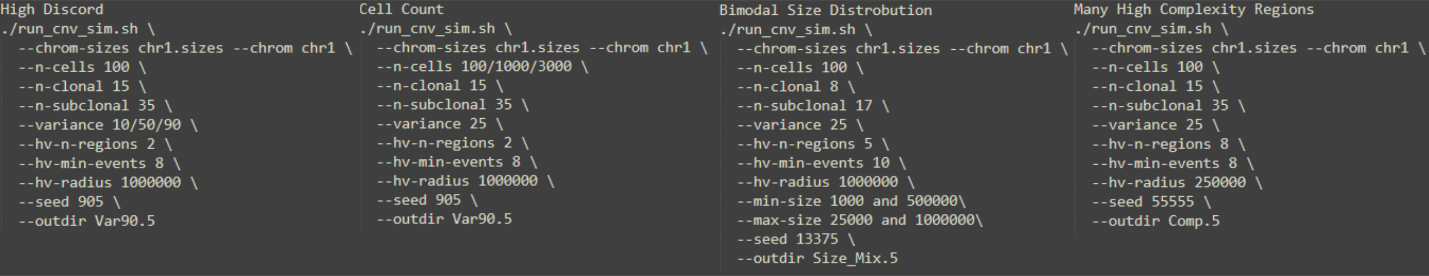

**Calculation of Jaccard Index and F1 score**

**Jaccard:** The Truth Set was filtered the intersection of the Truth Set and any possible overlaps across the sum total HG002 dataset. So, only those regions covered by any SVs in the sum total dataset were included as possible matches; conversely, no SVs were excluded from the sum total dataset. All SVs were maintained for two reasons 1) To further challenge the SV mergers in more complex regions, and 2) Maintain the diversity of SVs called by the SV callers.

bedtools jaccard was used to determine the similarity between files. It is based on base pair overlap, therefore, any differences are due to under/overmerging in regions where multiple alleles exist. Essentially, either under or over merging decrease overall similarity, even in regions were SVs overlap with the truth set.

- bedtools jaccard -a TruthSet.bed -b Merger.Bed > jaccard.txt

**F1 score:** The F1 score is calculate using the definitions of true positives, false positives, and false negatives as described in **figure 1** in the main text. True positives were determined by pure RO at three different stringencies. Essentially, post merged SVRs were compared back to the Truth Set and unique matches were identified. Non-unique matches to the Truth set were considered false negatives, while SVRs that did match the Truth Set were called as false positives (even if there were no possible SVs to match to in the Truth Set; in which all SV mergers should preform equally).

**True positives** (only returns unique matches to the Truth Set at a 50% RO)

- bedtools intersect -wa -r -u -f .5 -a TruthSet.ov.bed -b merger.sort.bed > ./ReRO.tp.bed
- #TP = n(ReRO.tp.bed)

**False negatives** (similar to True Positives, but non-unique matches are also included, the difference between these constitutes the number of false negatives)

- bedtools intersect -wa -r -f .5 -a TruthSet.ov.bed -b merger.sort.bed > ./ReRO.fn.bed
- #FN = n(ReRO.tp.bed) – n(ReRO.fn.bed)

**False positives** (were those that did not overlap at with the Truth Set at least a 50% RO)

- bedtools intersect -r -v -f .5 -a TruthSet.ov.bed -b merger.sort.bed > ./ReRO.fp.bed
- #FP = n(ReRO.fp.bed)

**SVR generation calculation**

Exact input parameters for SVCROWS used in the analysis of the application can be found in ***Table S3*.** An example of the input for both SVCROWS and CNVRuler are included in ***Table S1 & S2***. The general logic and formula that two SVs need to match reciprocally is:

[ (#Of_SV1_BP / #Of_SV2_BP) ≥ RO-Threshold && (#Of_SV2_BP / #Of_SV1_BP) ≥ RO-Threshold ]

**SV Merger usage**

**SVCROWS** Default was used for most analyses as it automatically adapts to datasets. Concise and Strict runs were adjusted based off of the parameters determined by the default run (xs ~= 1000, xl ~= 30000) to depict a range of stringencies without necessarily biasing the output. The SVCROWS algorithm can likely be more fine tuned than the three categories, but that was not the goal of the study. BPFactor and ExpandRORegion (see **Fig. S1**) were used to demonstrate their utility across the paper both for SV matching to a Truth Set, and SV Discovery.

- SVCROWSDefault (on all datasets)
  - Scavenge("~/R/SVCROWS/SVCROWSin", "~/R/SVCROWS/SVCROWSout", ExpandRORegion = TRUE, BPfactor = TRUE, DefaultSizes = TRUE)
    - In OC dataset smallSV = 1000bp, largeSV = 30000bp
    - In HG002 dataset smallSV = 332bp, largeSV = 818bp
      - Breakpoint matching boundries were 10% of those respective lengths
    - SVCROWSDefault is set at smallRO = 40%, largeRO = 70%
- SVCROWSConsice (on ovarian cancer dataset)
  - Scavenge("~/R/SVCROWS/SVCROWSin", "~/R/SVCROWS/SVCROWSout", ExpandRORegion = TRUE, BPfactor = TRUE, DefaultSizes = FALSE, xs = 1000, xl = 30000, y1s = 1000, y1l = 10000, y2s = 33, y2l = 66)
- SVCROWSStrict (on ovarian cancer dataset)
  - Scavenge("~/R/SVCROWS/SVCROWSin", "~/R/SVCROWS/SVCROWSout", ExpandRORegion = TRUE, BPfactor = TRUE, DefaultSizes = FALSE, xs = 1000, xs = 30000, y1s = 100, y1l = 1000, y2s = 66, y2l = 80)

**VCFTools** was used as a negative control merge; where only SVs with the same exact breakpoints and allele type were merged. For both datasets, all alleles were manually converted to <DUP> and <DEL>, and only those two alleles were used.

- vcf-merge A.vcf.gz B.vcf.gz C.vcf.gz … N.vcf.gz --collapse | bgzip -c > out.vcf.gz

**BCFTools** is used specifically for as recommended by the Truvari protocol. The recommended usage suggests to merge by ID, where all SVs have a unique ID. This method was used for the HG002 dataset, and yielded artifacts depicted in **Fig. S2B**. For the SC analysis, the merge key was changed to ALL, which helped reduce pure redundancy, and largely eliminated those artifacts.

- bcftools merge -m ID  file1.vcf.gz file2.vcf.gz file3.vcf.gz … filen.vcf.gz > out.vcf (HG002)
- bcftools merge -m ALL  file1.vcf.gz file2.vcf.gz file3.vcf.gz … filen.vcf.gz > out.vcf (SC)

**Truvari** uses a multifaceted algorithm to call overlaps, including an option for RO. We did not use the RO function to help distinguish the relative merits of different mergers; It would be more challenging to attribute SVCROWS’ performance to RO specifically. Further, Truvari tended to overmerge throughout this study, and the inclusion of the RO option (even at low percentages) either had no impact, or exacerbated the overmerging (data not shown).

- truvari collapse -k common –redist 1000 --chain --pctovl 0  -maxsize 100000000 -i bcftools.vcf.gz -o truvari.redux.out -c truvari.redux.collapsed.vcf

**Jasmine** also uses an algorithm that does not lend itself to a clear cause and effect in terms of its input and outputs. For similar reasons as Truvari, we did not include the RO option in usage of this program.

- jasmine file_list=filelist.txt max_dist_linear=0.7 min_overlap=0.05  out_file=merged.sizeweighted.vcf

**Svimmer** was run using 1000bp as limits for the two variables shown, in accordance with the other programs.

- ./svimmer --max_distance 1000 --max_size_difference 1000 oc.gz.txt 1 2 3 4 5 6 7 8 9 10 11 12 13 14 15 16 17 18 19 20 21 22 X Y > svimmer.out

**Survivor** is very similar to Svimmer’s algorithm, yet still distinct. 1000bp was still used as this study’s standard size difference.

- ./SURVIVOR merge path/to/files.txt [max size distance]1000 [one caller needed]1 [agree on type]1 [agree on strand]1 [GT]0 [simnimum size]10 survivor.out

**CNVRuler** has its own input format, similar to SVCROWS. It has 3 overlapping algorithms, but we elected to compare it against specifically its version of static RO. The input dataset can be found in **Table S1**.

**Links to data**

Links to the GIAB project data files:

HG002 SV Truth Set: <https://ftp-trace.ncbi.nlm.nih.gov/ReferenceSamples/giab/release/AshkenazimTrio/HG002_NA24385_son/NIST_SV_v0.6/>

MetaSV analysis of HG002 SV Truth Set: <https://ftp-trace.ncbi.nlm.nih.gov/ReferenceSamples/giab/data/AshkenazimTrio/analysis/BINA_Roche_MetaSV_10142016/HG002/>

LUMPY analysis of HG002 SV Truth Set: <https://ftp-trace.ncbi.nlm.nih.gov/ReferenceSamples/giab/data/AshkenazimTrio/analysis/BU_GRCh38_SVs_06252018/>

mrCarNaVar analysis of HG002 SV Truth Set: <https://ftp-trace.ncbi.nlm.nih.gov/ReferenceSamples/giab/data/AshkenazimTrio/analysis/BilkentUni_IlluminaHiSeq_VariationHunter_mrCaNaVar_08282015/>

Sniffles analysis of HG002 SV Truth Set: <https://ftp-trace.ncbi.nlm.nih.gov/ReferenceSamples/giab/data/AshkenazimTrio/analysis/Baylor_sniffles_05092017/>

Pbsv analysis of HG002 SV Truth Set: <https://ftp-trace.ncbi.nlm.nih.gov/ReferenceSamples/giab/data/AshkenazimTrio/analysis/PacBio_HiFi-Revio_20231031/pacbio-wgs-wdl_germline_20231031/>

Link to EGA data (SC dataset):

<https://ega-archive.org/datasets/EGAD00001009456>
